# Propagating actomyosin-generated force to intercellular junction

**DOI:** 10.1101/191965

**Authors:** Vivian W. Tang

**Affiliations:** Department of Cell and Developmental Biology, University of Illinois, Urbana-Champaign

**Author notes:** Corresponding author Address correspondence to: Vivian W. Tang, Department of Cell and Developmental Biology, University of Illinois, Urbana-Champaign, 601 S. Goodwin Avenue, Urbana, IL 61801.

**Keywords:** alpha-actinin, synaptopodin, myosin, junction, mechanobiology, cell-cell

## Abstract

Actomyosin II contractility in epithelial cells plays an essential role in tension-dependent adhesion strengthening. One key unsettling question is how cellular contraction transmits force to nascent cell-cell adhesion when there is no stable attachment between the nascent adhesion complex and actin filament. Here, we showed that application of intercellular tension induces myosin 1c accumulation at the lateral membrane between epithelial cells. We hypothesized that the accumulation of myosin 1c at the cell-cell interface allows coupling of actomyosin contractility to the generation of intercellular tension, thus is essential for tension-induced junction maturation. We showed that myosin 1c KD compromises a-actinin-4 recruitment to cell-cell adhesion during normal junction maturation driven by endogenous actomyosin contractility. However, application of cyclic tension to intercellular junction from outside of the cells rescued tension-dependent a-actinin-4 accumulation, suggesting that myosin-1c KD did not compromise the tension response or disrupt the overall integrity of the junctional complex. Our study identifies myosin-1c as a novel tension-sensitive protein on the lateral membrane and underscores a non-junctional contribution to adhesion strengthening at the epithelial cell-cell adhesion interface.

## Introduction

Epithelial cells are frequently subjected to chemical and mechanical insults, leading to a local loss of cells from the epithelium. In order to restore a continuous barrier, epithelial cells must quickly migrate into the open wound and reseal the monolayer [1-5]. However, covering the wounded area is not sufficient for the new epithelium to provide resilience at the tissue-level. The regeneration process must create mature junctions with strong cell-cell adhesion and attachment to the actin cytoskeleton. Failure to strengthen cell-cell adhesion and re-establish mature junctions upon wound closure predisposes the epithelium to future injuries, resulting in chronic conditions [4, 6-9]. Understanding how a confluent epithelial monolayer strengthen cell-cell adhesion can provide insight into the final stages of epithelial regeneration and help identify ways to facilitate reformation of a strong epithelial protective layer [10-12].

Junction formation starts with a nucleation process initiated by trans-interaction of the cadherin-catenin complex from apposing cells [13, 14], followed by a maturation process that requires mechanical input from actomyosin contractility [15, 16], leading to stabilization of the cadherin-catenin complex [17-20], recruitment of additional junctional molecules [15, 21-27], and attachment to actin filaments [28-33]. Adhesion strengthening in an epithelial cell sheet is governed partly by tension-induced recruitment of *α*-actinin-4 to cell-cell interactions in a synaptopodin-dependent manner [21]. Yet, it remains unclear how actomyosin contractility propagate mechanical force to generate tension at cell-cell adhesions. Synaptopodin forms a complex with myosin II and could potentially link actomyosin contractility to adhesion molecules [21]. However, synaptopodin recruitment to the cell junction is dependent on junctional tension. Therefore, an upstream mechanism must exist to create tension at cell-cell adhesion prior to the recruitment of synaptopodin and *α*-actinin-4. Here, we have identified an upstream regulator of *α*-actinin-4 as the membrane-associated actin-binding motor, myosin-1c [34-41].

## Results

### Myosin-1c colocalizes with α-actinin-4 and synaptopodin on the lateral membrane

To identify potential regulator of *α*-actinin-4 on the plasma membrane, we used a proximity crosslinking approach and identified myosin-1c in mature junctional complexes (Supplementary Fig. 1, see Methods). Myosin-1c has previously been shown to regulate E-cadherin adhesion, thus could potential participate in junction maturation through the regulation of *α*-actinin-4. To characterize the relationship between myosin-1c and *α*-actinin-4, we performed immunofluorescence localization using optical-sectioning structured illumination microscopy (OS-SIM). In polarized epithelial cells, myosin-1c and *α*-actinin-4 show significant correlation on the linear junction (Fig. 1A-B, oblong and Supplementary Fig. 2). At the vertices of linear junctions, myosin-1c shows correlation with both synaptopodin and *α*-actinin-4 (Fig. 1B, circle and Supplementary Fig. 2). Furthermore, myosin-1c overlaps with E-cadherin and *α*-catenin at discreet regions along the lateral junctions (Supplementary Fig. 3-4). On the contrary, myosin IIB preferentially accumulates at the apical junction but is mostly absent on the lateral adhesions (Supplementary Fig. 5-7). These observations indicate that myosin-1c is a major component on the lateral membrane which co-localizes with junctional complexes and tension-sensitive molecules, suggesting a possible role of myosin-1c in tension-dependent regulation of cell-cell adhesion on the lateral interface.

**Figure 1.**
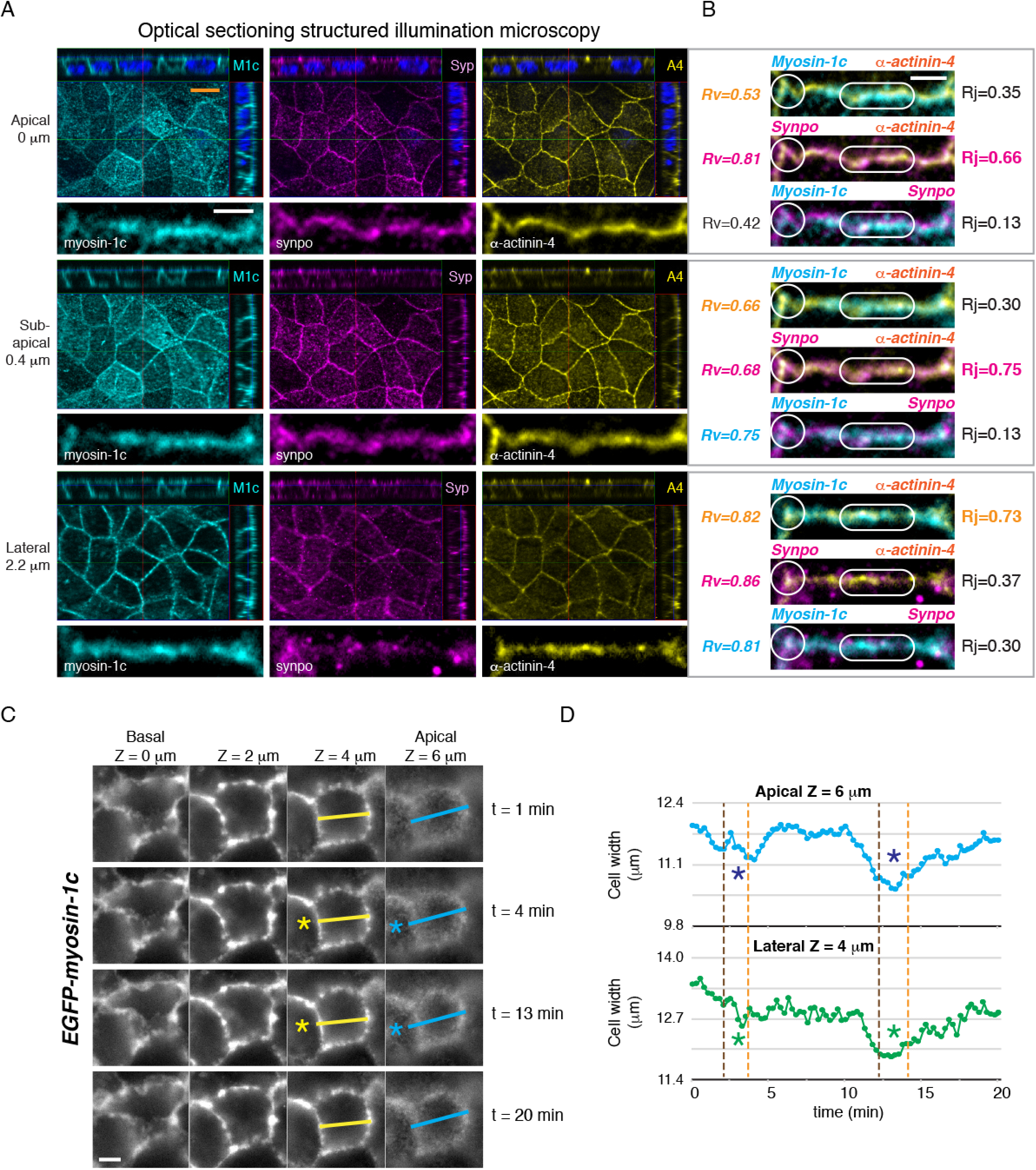
Myosin-1c is associated with tension-sensitive junctional complex and tethered on the lateral membrane during whole cell contraction in polarized epithelial monolayer. (A) Optical sectioning structured illumination microscopy showing immunofluorescence localization of myosin-1c (M1c), synaptopodin (Syp), and *α*-actinin-4 (A4) at the apical junction (Z=0 μm), sub-apical junction (Z=0.4 μm), and lateral junction (Z=2.2 μm). For each Z-image, the top panel shows the X-Z, Y-Z, and X-Y images and the bottom panel shows a representative linear junction from the X-Y-image. Data is representative of 10 sets of images (N=10) from one experiment out of 6 independent experiments (N=6). Orange scale bar is 10 μm and white scale bar is 2 μm. (B) Merge images of the representative linear junction shown in A. Pearson’s correlation coefficient between myoin-1c, synaptopodin (synpo), or *α*-actinin-4 at junctional vertices (blue circle) is represented by Rv and at linear junction (blue oblong) is represented by Rj. Correlations were calculated from 500-1500 pixels per protein per z-image per one set of data (see Figure S2). Six sets of data (N=6) were analyzed and a representative set of data is shown. Scale bar is 2 μm. (C) Time-lapse frames from Movie 1 showing EGFP-myosin-1c along the entire lateral cell-cell interface (from apical Z=6 μm to basal Z=0 μm) during a single whole contraction (yellow and blue asterisks). Data is representative of 8 sets of live-cell time-lapse movies (N=8). Scale bar is 5 μm. (D) Measurements of cell width (marked by yellow and blue lines in C) and EGFP-myosin-1c intensity during whole contraction (Movie 1).

**Figure 2.**
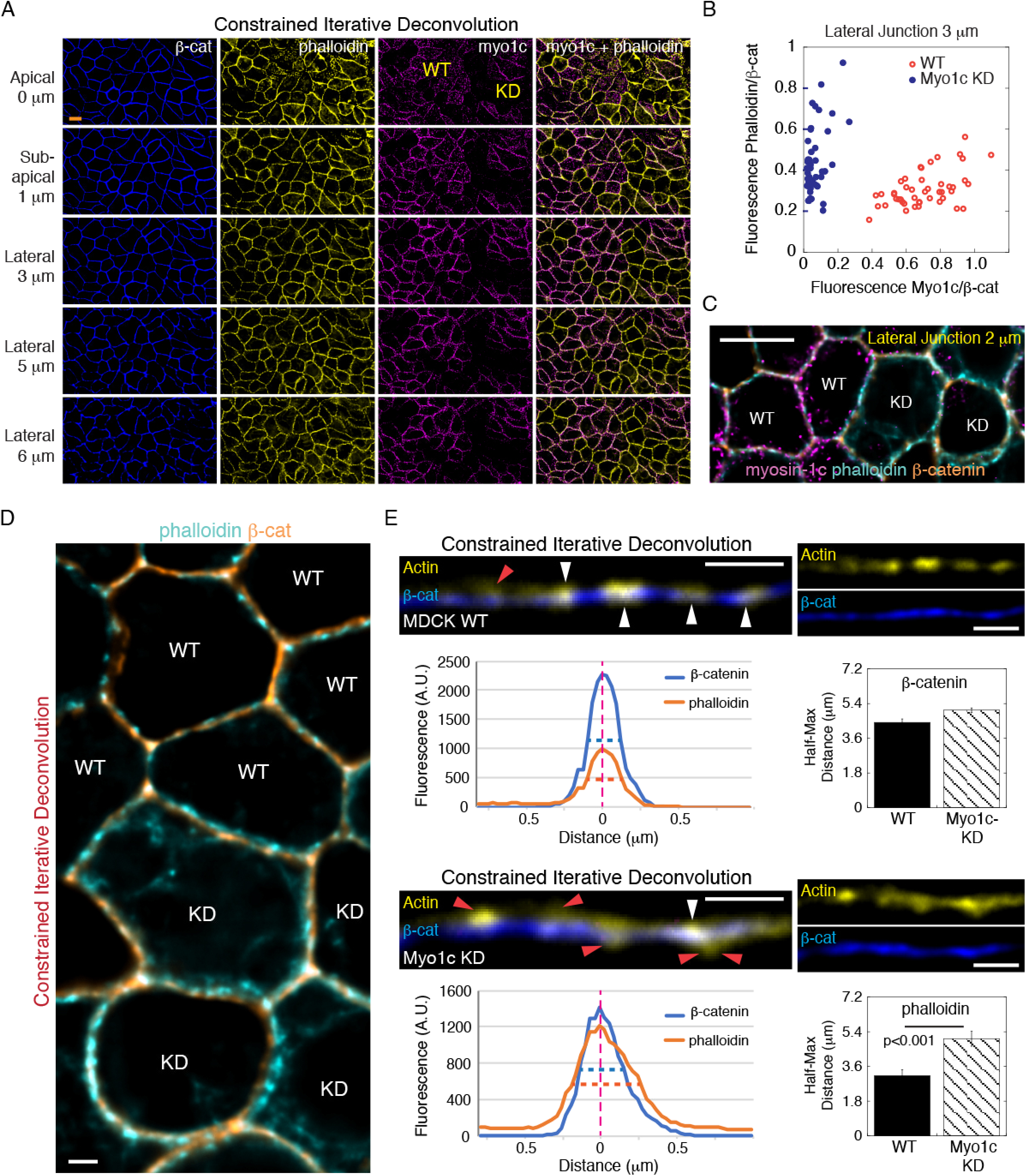
Myosin-1c knockdown disrupts actin compaction on the lateral plasma membrane. (A) Constrained iterative deconvolution microscopy showing immunofluorescence localization of myosin-1c, actin (phalloidin), and *β*-catenin at the apical junction (Z=0 μm), sub-apical junction (Z=1 μm), and lateral (Z=3, 5 and 5μm) junctions. Images were generated from 0.2 μm wide-field optical sections. Wild-type and myosin-1c knockdown cell areas are marked by WT and KD, respectively. Data is representative of 10 sets of images (N=10) from one experiment out of 3 different experiments (N=3). Orange scale bar is 10 μm and white scale bar is 2 μm. (B) Junctional level of myosin-1c and actin (phalloidin) in wild-type and myosin-1c knockdown cells. Each point represents measurement of one junctional area at Z=3 μm. Data is representative of 10 sets of images (N=10) from one experiment out of 3 independent experiments (N=3). (C) Merge image of myosin-1c, actin (phalloidin), and *β*-catenin at the lateral junction (Z=2 μm) from A. Wild-type and myosin-1c knockdown cell areas are marked by WT and KD, respectively. Scale bar is 10 μm. (D) Merge image of actin (phalloidin) and *β*-catenin at the lateral junction (Z=2 μm) from A. Wild-type and myosin-1c knockdown cell areas are marked by WT and KD, respectively. Scale bar is 2 μm. (E) Actin dissociates from *β*-catenin at the lateral junction (Z=2 μm) in myosin-1c knockdown cells. Images of actin (phalloidin) and *β*-catenin (*β*-cat) at the lateral junction (Z=2 μm) in wild-type (WT) and myosin-1c knockdown (Myo1c KD) cells from A. White arrowheads mark junctional regions with actin (phalloidin) overlapping with *β*-catenin. Orange arrowheads mark junctional region with actin dissociated from *β*-catenin. Line graphs show intensities of actin (phalloidin) and *β*-catenin across the lateral junctions. Bar graphs show the mean and standard errors of the half-maximal width of actin (phalloidin) and *β*-catenin across 12 junctions (N=12). Data is representative of 10 sets of images (N=10) from one experiment out of 3 independent experiments (N=3). Scale bar is 2 μm.

Using live-cell imaging of EGFP-myosin-1c, we found that myosin-1c is tethered on the lateral plasma membrane during endogenous cellular contractions (Fig. 1C, yellow and blue asterisks and movies 1-2). Measurements of cell width show decreases in cell width along the entire lateral cell-cell interface (Fig. 1D), indicating that contraction is associated with constriction on the X-Y cross-section plane of the cell. Since EGFP-myosin-1c is stably tethered on the plasma membrane during cellular contractions, it can potentially play a role in force transmission by coupling actomyosin II contractility to the cell membrane.

### Myosin-1c knockdown disrupts actin compaction on the lateral membrane

To examine the role of myosin-1c in the organization of actin on the lateral cell-cell interface, we depleted myosin-1c in MDCK cells using shRNA. Myosin-1c knockdown significantly disrupts actin organization on the lateral membrane without changing the level of actin and *β*-catenin (Fig. 2A-D and Supplementary Fig. 8). Myosin-1c and actin usually overlap with *β*-catenin at discreet regions on the lateral junctions (Fig. 2E, white arrowheads). However, in myosin-1c knockdown cells, actin and *β*-catenin are spatial separated (Fig. 2E, red arrowheads), indicating a defect in the compaction of cortical actin. Measurement of actin intensity tangential to the junction shows a ∼1.8-fold increase (from ∼0.36 μm to ∼0.68 μm) in the half-maximum thickness of cortical actin in myosin-1c knockdown cells. These results show that myosin-1c is important for coupling actin and the cadherin-catenin complex.

### Myosin-1c knockdown compromises coupling between actomyosin II contractility and the lateral membrane

Cellular contractility can be globally regulated by intracellular calcium. To further characterize the changes in cell width during whole cell contraction, intracellular calcium and membrane movement were simultaneously monitored using a lipid-modifiable calcium-sensitive intramolecular FRET sensor Lyn-D3cpV (Supplementary Figure 9 and Movies 3-5). Live-cell imaging of Lyn-D3cpV in young monolayer shows that whole cell contraction is preceded by a transient rise of intracellular calcium followed by a decrease in cell width (Supplementary Figure 9A and Movies 3-4). In maturing cell monolayers, a cell can be stretched by the contraction of its neighbor (Supplementary Figure 9B and Movie 5). Prolonged stretch at a junctional area (Supplementary Figure 9B, blue arrowhead) can lead to increase in cytoplasmic calcium in the cell that is being stretch (Supplementary Figure 9B, blue arrowhead), resulting in a cellular contraction and a counter-tug at the neighboring cell (Movie 5). These observations indicate that whole cell contraction could potentially create tension at cell junction if force can be transmitted from the contracting cytoskeleton to the junctional complex on the plasma membrane.

Previously, we showed that exogenous application of intercellular force induces recruitment of a-actinin-4 during junction maturation [21]. Here, we found that endogenous cellular contraction can elicit local recruitment of venus-*α*-actinin (Supplementary Fig. 10A-C, blue circle and line and Movie 6). A non-contracting junction just a few microns away in the same cell does not recruit venus-*α*-actinin (Supplementary Fig. 10A-C, orange circle and line and Movie 6). These observations indicate that a single and spatially restricted contraction is sufficient to elicit molecular change(s) locally at the adhesion complex, underscoring a mechanism that uses actomyosin II contractility to recruit *α*-actinin to the junction in a tension-dependent manner.

In young and maturing junction, repeated whole cell contraction increases venus-*α*-actinin ∼20% in less than 5 hours (Supplementary Fig. 10D-F and Movie 7). Contractions in young and maturing monolayers occurs along the entire lateral membrane (Supplementary Fig. 10G, and Movies 8-12). Contraction in young monolayer is associated with gradual accumulation of venus-*α*-actinin at *α*-actinin-negative junction (Supplementary Fig. 10H) and transient accumulation of venus-*α*-actinin on the nuclear envelope (Supplementary Fig. 10I-J, Supplementary Fig. 11, and Movie 8-12). These observations implicate that, in young epithelial monolayer, a mechanism of force transmission must exist to propagate actomyosin II contractions to cell-cell adhesions prior to junction maturation and stable linkage of actin to adhesion complexes.

To examine the role of myosin-1c in tension-induced *α*-actinin-4 recruitment, we depleted myosin-1c in venus-*α*-actinin expressing cells. We found that knockdown of myosin-1c compromises venus-*α*-actinin localization to cell-cell interface and alters the behavior of cellular contractions (Fig. 3A-C and Movie 13-14). Moreover, contractions of myosin-1c KD cells did not result in venus-*α*-actinin accumulation at the cell junction despite normal translocation onto the nuclear envelop (Fig. 3A-B). Instead of a reduction in cell width during contraction, myosin-1c knockdown cell expands, resulting in an increase in the cell area (Fig. 3C and Movie 13-14). Furthermore, myosin-1c knockdown cells constantly blebs at the lateral cell-cell interface (Movies 15-16), a phenomenon that happens rarely in wild-type cells (Movie 17). These observations indicate that the integrity of the lateral membrane in myosin-1c knockdown cells is compromised.

**Figure 3.**
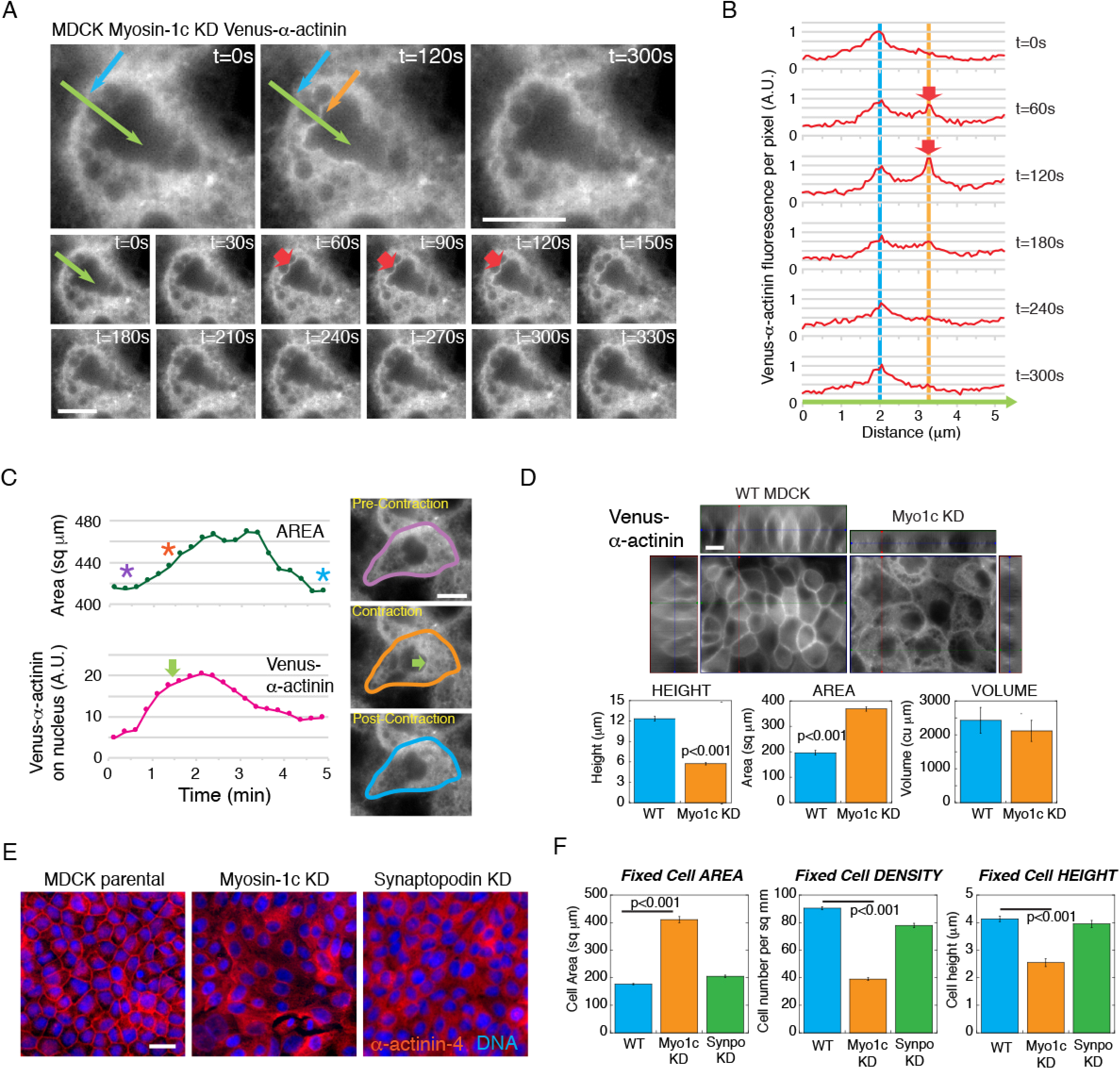
Myosin-1c knockdown compromises the coupling between actomyosin contractility and the lateral plasma membrane. (A) Time-lapse frames from Movie 13 showing a lack of junction accumulation of venus-*α*-actinin (blue arrow) during or after contraction despite transient accumulation on the nuclear envelope during contraction (t=60-150s, orange and red arrows). Green arrow marks the line-scan used to generate graph in B. Data is representative of 6 sets of live-cell time-lapse movies (N=6). (B) Measurements of venus-*α*-actinin intensities along the green arrow in A. Blue and orange dotted lines mark the cell junction and nuclear envelope, respectively. Red arrows point to increase in venus-*α*-actinin intensities on the nuclear envelope during contraction (t=60 and 120s). (C) Measurement of cell area and venus-*α*-actinin intensities on the nuclear envelope during contraction (t=1.5 min, orange asterisk and green arrow) of the cell in D and Movie 14. Purple and blue asterisks mark pre- and post-contraction, respectively. Time-lapse frames from movie 14 showing areas representing pre-contraction (purple outline), contraction (orange outline) and post-contraction (blue outline). Green arrow points to the transient accumulation of venus-*α*-actinin on the nuclear envelope during contraction. Data is representative of 6 sets of live-cell time-lapse movies (N=6). (D) Live-cell images of mature monolayers of wild-type (WT) and myosin-1c (Myo1c) knockdown (KD) cells expressing venus-*α*-actinin from Movies 16 and 17. X-Z, Y-Z, and X-Y images are used to calculate cell height, spread area, and cell volume in G. Image set is representative of 16 sets of images (N=16) from one experiment out of 4 independent experiments (N=4). Scale bars are 10 μm. Bar graphs show measurements of cell height, spread area, and cell volume of wild-type (WT) and myosin-1c (Myo1c) knockdown cells. Mean and standard errors of 16 measurements (N=16) are plotted. Data is representative of one experiment out of 4 independent experiments (N=4). (E) Widefield immunofluorescence images of *α*-actinin-4 in wild-type and myosin-1c knockdown cell monolayers. DNA is stained with Hoechst. Image set is representative of 16 sets of images (N=16) from one experiment out of 3 independent experiments (N=3). Scale bar is 10 μm. (F) Monolayer spread area, cell density, and cell height of wild-type (WT), myosin-1c knockdown (Myo1c KD), and synaptopodin knockdown (Synpo KD) cells. Bar graphs show the means and standard errors of 16 measurements (N=16). Data is representative of one experiment out of 4 independent experiments (N=4).

Knockdown of myosin-1c diminishes the formation of the lateral membrane in venus-*α*-actinin expressing cells (Fig. 3D). In fixed cells, knockdown of myosin-1c abolished junctional localization of endogenous *α*-actinin-4 (Fig. 4E). The inability for myosin-1c knockdown cells to develop proper cell height is not due to a lack of *α*-actinin-4 at the junction since synaptopodin knockdown cells that are also defective in junctional recruitment of *α*-actinin-4 developed normal cell height (Fig. 3E-F). These findings indicate that myosin-1c plays an important role in the generation of the lateral membrane and is essential for the accumulation of *α*-actinin at cell junction.

**Figure 4.**
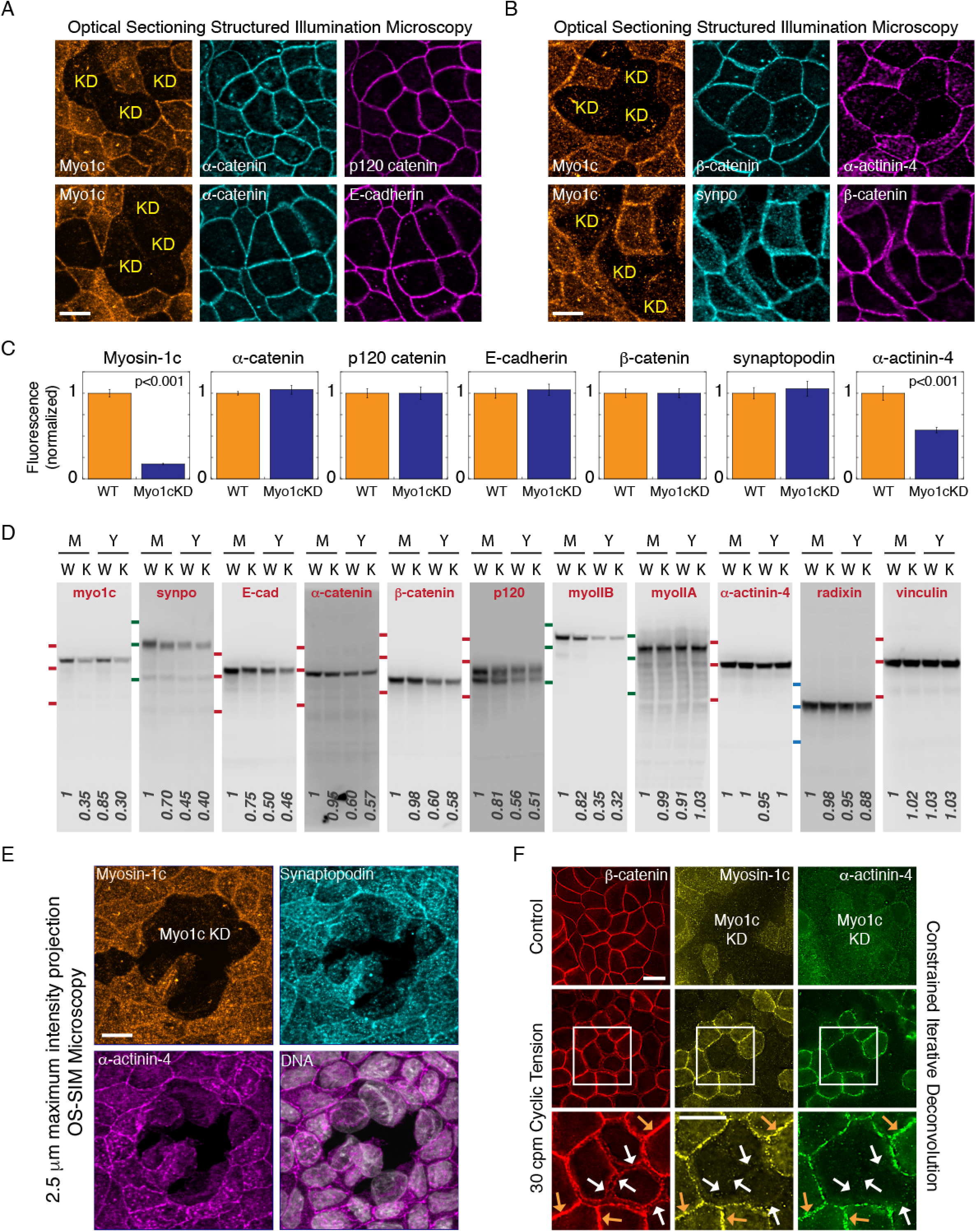
Myosin-1c knockdown abolishes junctional recruitment of *α*-actinin-4 and weakens cell-cell cohesion. (A) OS-SIM images of myosin-1c (Myo1c), *α*-catenin, p120-catenin, and E-cadherin at the apical plane of heterogeneous myosin-1c knockdown (KD) cell monolayer. Data is representative of 12 sets of images (N=12) from one experiment out of 3 independent experiments (N=3). (b) OS-SIM images of myosin-1c (Myo1c), *β*-catenin, synaptopodin (synpo), and *α*-actinin-4 at the apical plane of heterogeneous myosin-1c knockdown (KD) cell monolayer. Data is representative of 12 sets of images (N=12) from one experiment out of 3 different independent (N=3). (C) Quantification of junctional myosin-1c, *α*-catenin, p120-catenin, E-cadherin, *β*-catenin, synaptopodin, and *α*-actinin-4 intensities in wild-type and myosin-1c knockdown cells. Bar graphs show mean intensity and standard errors of 24 junctions (N=24). Data is representative of one experiment out of 3 independent experiments (N=3). (D) Western blots for myosin-1c (myo1c), synaptopodin (synpo), E-cadherin (E-cad), *α*-catenin, *β*-catenin, p120-catenin, myosin IIB (myoIIB), myosin IIA (myoIIA), *α*-actinin-4, radixin, and vinculin in cell lysates from mature (M) and young (Y) confluent monolayers of wild-type (W) and myosin-1c knockdown (K) cells. Data is representative of one set of experiments out of 4 independent experiments (N=4). Red MW markers are 135, 100, 75 kD. Green MW markers are 180, 135, 100 kD. Blue MW markers are 100, 75, 50 kD. (E) Maximum intensity projections from a 2.5 μm z-stack of 0.2 μm OS-SIM z-images showing immunofluorescence of myosin-1c, synaptopodin, and *α*-actinin-4 in mature cell monolayer with heterogeneously knockdown of myosin-1c (Myo1c KD). Image set is representative of 12 image sets (N=12) from one experiment out of 6 independent experiments (N=6). Scale bar is 10 μm. (F) Constrained iterative deconvolution microscopy showing immunofluorescence of myosin-1c, *α*-actinin-4, and *β*-catenin in a young monolayer with heterogeneous myosin-1c knockdown (Myo1c KD) that has been treated or untreated (control) with cyclic tension. Image set is representative of 8 image sets (N=8) from one experiment out of 4 independent experiments (N=4). Scale bar is 10 μm.

### Myosin-1c knockdown abolishes junctional recruitment of α-actinin-4

To investigate the relationship between myosin-1c and junctional proteins, we compared their localization and total cellular levels in confluent monolayers of wild-type and myosin-1c knockdown cells. However, as shown in Fig. 3D-F, myosin-1c knockdown cells are shorter, which makes comparison between wild-type and myosin-1c knockdown cells problematic due to a difference in their Z-height. In order to make a fair assessment, we standardized a protocol that forces myosin-1c knockdown cells to exhibit a lateral membrane similar to the wild-type cells. When myosin-1c knockdown cells are plated onto a Transwell insert/cup at 100% confluent density (that of wild-type cells), they are forced to squeeze among themselves to create a lateral membrane. Using this protocol, we found that none of the canonical adhesion proteins, E-cadherin, *α*-catenin, *β*-catenin, or p120-catenin, is affected by myosin-1c knockdown (Fig. 4A-D).

OS-SIM projections and quantitation of junctional intensity show that only *α*-actinin-4 is reduced in myosin-1c knockdown cells (Fig. 4A-C). The junctional levels and the localization of E-cadherin, *α*-catenin, *β*-catenin, p120-catenin, and synaptopodin were not affected by knockdown of myosin-1c. However, myosin-1c does have an indirect role at the junction. Usually, upon confluency, adhesion complexes continue to mature and the levels of E-cadherin, *α*-catenin, *β*-catenin, p120-catenin, synaptopodin, and myosin IIB would increase over several days (Fig. 4D, young, Y and mature, M). Knockdown of myosin-1c slightly retards the increase of E-cadherin and synaptopodin over this maturation period (Fig. 4D). Thus, myosin-1c does not considerably affect the expression of junctional proteins but plays an indirect role in their accumulation over time during adhesion strengthening and epithelial maturation.

### Exogenous Tension rescues junctional recruitment of α-actinin-4 in myosin-1c knockdown cells

The lack of junctional *α*-actinin-4 in myosin-1c knockdown cells may due to defective force transmission. If the problem of *α*-actinin-4 localization in myosin-1c knockdown cells is purely biophysical in nature, *α*-actinin-4 might still respond to exogenously-applied tension and recruit to the junction. We have previously developed a method to create tension at cell-cell adhesion in a monolayer of epithelial cells [21]. Using this method, pulsatile and cyclic force were applied to myosin-1c-knockdown cells. Usually, MDCK monolayer can withstand a medium level of cyclic force (∼20-100 nN per cell) and respond by strengthening their cell junction [21]. However, myosin-1c knockdown cells are more prone to spontaneous tearing when embedded in wild-type cells (Fig. 4E) and have altered response to exogenously applied force (Fig. 4F). We found that application of medium level of cyclic force (∼20 nN per cell at 30 cycles per min for 10 min) failed to induce junctional recruitment of synaptopodin and *α*-actinin-4 in myosin-1c knockdown cells. Instead, the applied force causes cell-cell interactions to break open at myosin-1c and *α*-actinin-4-negative junctions (Fig. 4F, white arrows). These findings show that myosin-1c-dependent recruitment of junctional *α*-actinin-4 is essential for strong cohesion among epithelial cells.

In order to test whether myosin-1c knockdown cells retains their ability to recruit a-actinin-4, we decreased the amount of force applied to the junction. Application of low level of pulsatile force (∼7 nN per cell at 8 cycles per min for 90 minutes) was able to trigger a full recovery of *α*-actinin-4 at the cell junction (Fig. 5A-B). These observations indicate that the molecular players and mechanisms that support tension-dependent junction maturation is completely intact in myosin-1c knockdown cells. After monolayer maturation, cyclic force (∼10 nN per cell at 20 cycles per min for 60 minutes) can induce an increase in junctional *α*-actinin-4 transiently above the steady-state level of a mature junction (Fig. 5C). Upon tension release, the level of junctional *α*-actinin-4 returns to a pre-tension steady-state level of a mature monolayer. In mature myosin-1c knockdown cells, the level of junctional *α*-actinin-4 is at a lower steady-state (Fig. 5C). Cyclic force (∼10 nN per cell at 20 cycles per min for 60 minutes) induced an increase in *α*-actinin-4 at myosin-1c-depleted junction to the same level as wild-type junctions. Upon tension release in myosin-1c knockdown cells, junctional *α*-actinin-4 is reset to a new higher level similar to the steady-state found in mature monolayer of wild-type cells. These results show that the retention of *α*-actinin-4 at cell junction does not require the continual presence of myosin-1c on the lateral membrane. Therefore, the dependency of *α*-actinin-4 on myosin-1c is biophysical in nature and restricted to the early stages of adhesion maturation during tension-driven remodeling.

**Figure 5.**
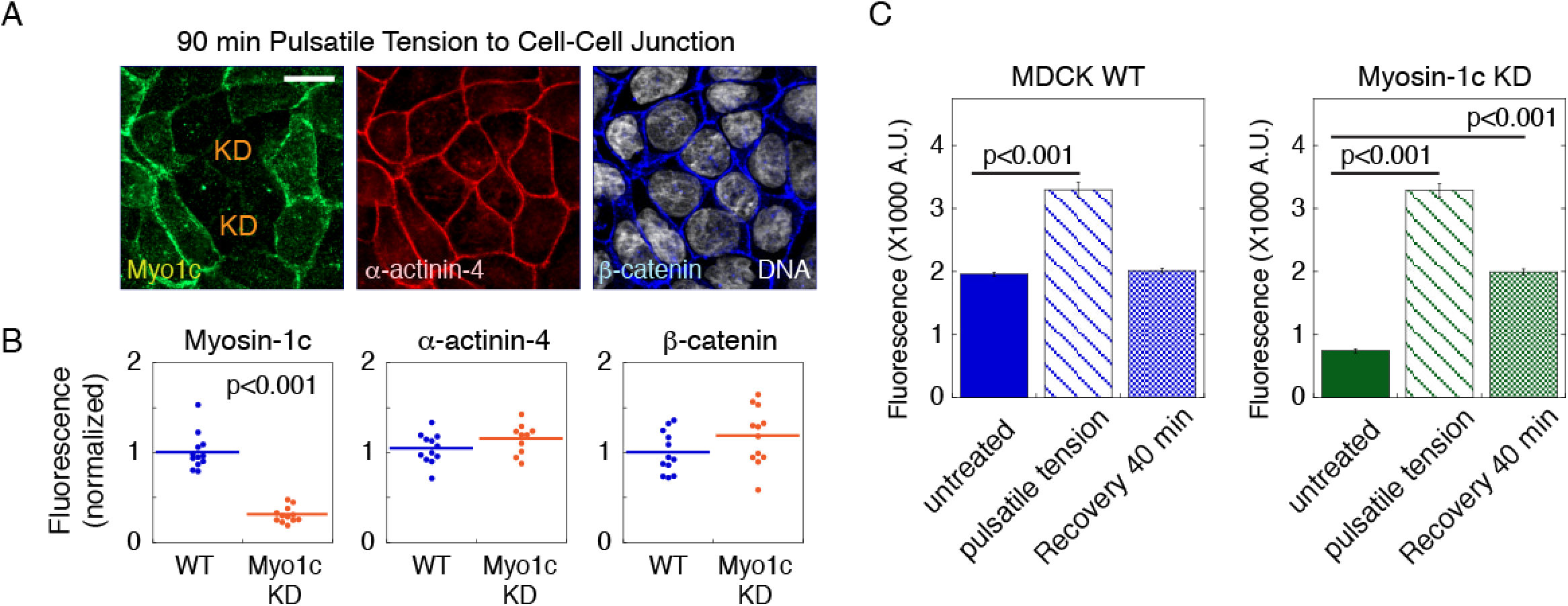
Junctional localization of *α*-actinin-4 can be rescued by exogenously applied tension in myosin-1c knockdown cells. (A) Maximum intensity projections from a 2.5 μm z-stack of 0.2 μm OS-SIM z-images showing myosin-1c, *α*-actinin-4, and *β*-catenin at the apical junction of monolayers treated with 90 min pulsatile tension. DNA is stained with Hoechst. Data is representative of one experiment out of 6 independent experiments (N=6). (B) Quantification of junctional myosin-1c (Myo1c), *α*-actinin-4, and *β*-catenin immunofluorescence intensities in wild-type (WT) and myosin-1c knockdown (Myo1c KD) monolayers treated with 90 min pulsatile tension. Horizontal lines represent mean fluorescence intensities. Data is representative measurements from one experiment out of 6 independent experiments (N=6). Scale bars are 10 μm. (C) Quantification of junctional *α*-actinin-4 immunofluorescence intensity in wild-type (WT) and myosin-1c knockdown (Myo1c KD) monolayers before (untreated), immediately after (pulsatile tension), and 40 mins after recovery from (recovery 40 min) treatment with 90 min pulsatile tension. Bar graphs show the mean and standard errors of 36 junctions (N=36). Data is representative of one experiment out of 3 independent experiments (N=3).

### Myosin-1c is recruited to the lateral membrane in a tension-sensitive manner

To further investigate the regulation of myosin-1c by intercellular tension, we applied low level of cyclic force (∼7 nN per cell at 20 cycles per min for 60 minutes) to young monolayers (Fig. 6A). Quantitation of z-images shows that myosin-1c and *α*-actinin-4 are induced at the apical and lateral junctions (Fig. 6B-C) whereas *β*-catenin exhibits only a minor change (Fig. 6D). Importantly, myosin-1c is upregulated to a greater extent on the lateral junctions than at the apical junction (Fig. 6B, 2-6 μm z-focal plane). On the contrary, *α*-actinin-4 is upregulated to a greater extent at the apical junction than at the lateral junctions (Fig. 6C, 1 μm z-focal plane). After cyclic force (∼7 nN per cell at 20 cycles per min for 60 minutes), junctional *α*-actinin-4 resets to a new higher steady-state in ∼15 min whereas membrane myosin-1c takes ∼30 min to reach its new steady-state (Fig. 6E-G). These results suggest that the mechanism that regulates tension-induced myosin-1c accumulation is different than that of *α*-actinin-4.

**Figure 6.**
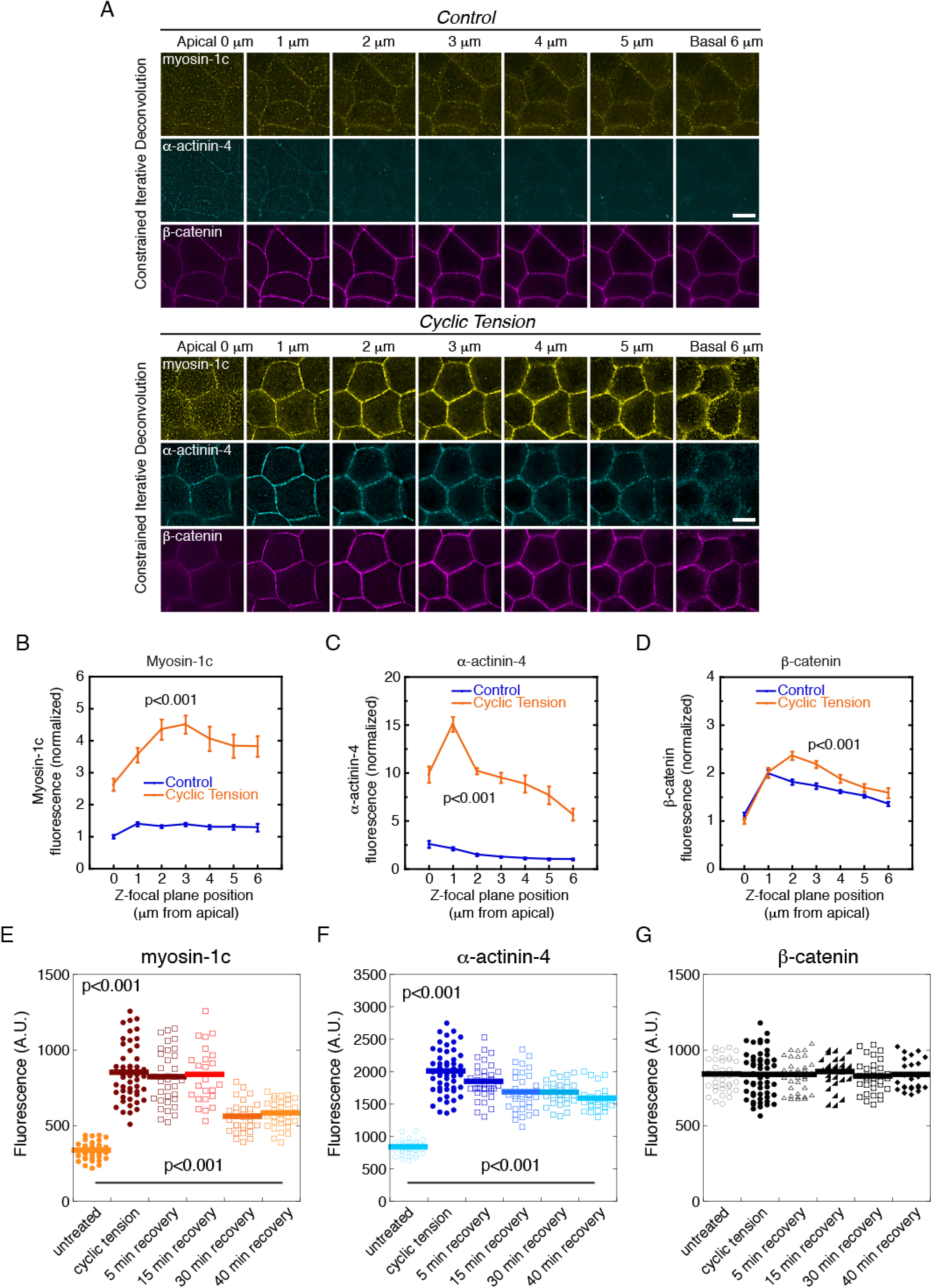
Myosin-1c is recruited to the lateral membrane in a tension-sensitive manner. (A) Deconvolved Z-slice images of myosin-1c, *α*-actinin-4, and *β*-catenin immunofluorescence in control or cyclic tension-treated polarized epithelial monolayers. Images were generated from constrained iterative deconvolution from 0.2 μm wide-field optical sections (from apical Z= 0 μm to basal Z= 6 μm). Data is representative of 12 sets of images (N=12) from one experiment out of 4 independent experiments (N=4). Scale bars are 10 μm. (B-D) Quantification of myosin-1c, *α*-actinin-4, and *β*-catenin intensities from apical (Z= 0 μm) to basal (Z= 6 μm) junctions in control and cyclic tension treated monolayers. Line graphs show the mean and standard errors of 24 junctions (N=24). Data is representative of one experiment out of 5 independent experiments (N=5). (E-G) Quantification of myosin-1c, *α*-actinin-4, and *β*-catenin immunofluorescence intensities at the apical junction in polarized cell monolayers before (untreated), immediately after (cyclic tension), and 5, 15, 30, 40 mins after recovery from treatment with 60 min cyclic tension. Data is representative measurements from one experiment out of 3 independent experiments (N=3).

To understand the relationship between synaptopodin, *α*-actinin-4, and myosin-1c, we compared their junction localization and intensity before and after force application. Exogenously cyclic force (∼10 nN per cell at 20 cycles per min for 60 minutes) increases the levels of junction myosin-1c, synaptopodin, and *α*-actinin-4 (Fig. 7A, white arrows) which show strong correlation with each other after tension (Fig. 7B-C). On the contrary, *β*-catenin shows no correlation with myosin-1c upon force application (Fig. 7D). These results implicate a possible tension-dependent relationship between synaptopodin, *α*-actinin-4, and myosin-1c. However, using synaptopodin and *α*-actinin-4 knockdown cells, we found that the tension response of myosin-1c is completely unaltered in the absence of synaptopodin or *α*-actinin-4 (Fig. 7E-H). Furthermore, in mature junction, a high level of cyclic force (∼50 nN per cell at 30 cycles per min for 60 min) can induce additional myosin-1c recruitment to the cell junction without eliciting a synaptopodin response (Fig. 7I-J), indicating that the tension response of myosin-1c continues to exist after junction maturation. These observations indicate that the mechanism that regulates the tension response of myosin-1c is independent of the mechanism that regulates the tension response of synaptopodin and *α*-actinin-4.

**Figure 7.**
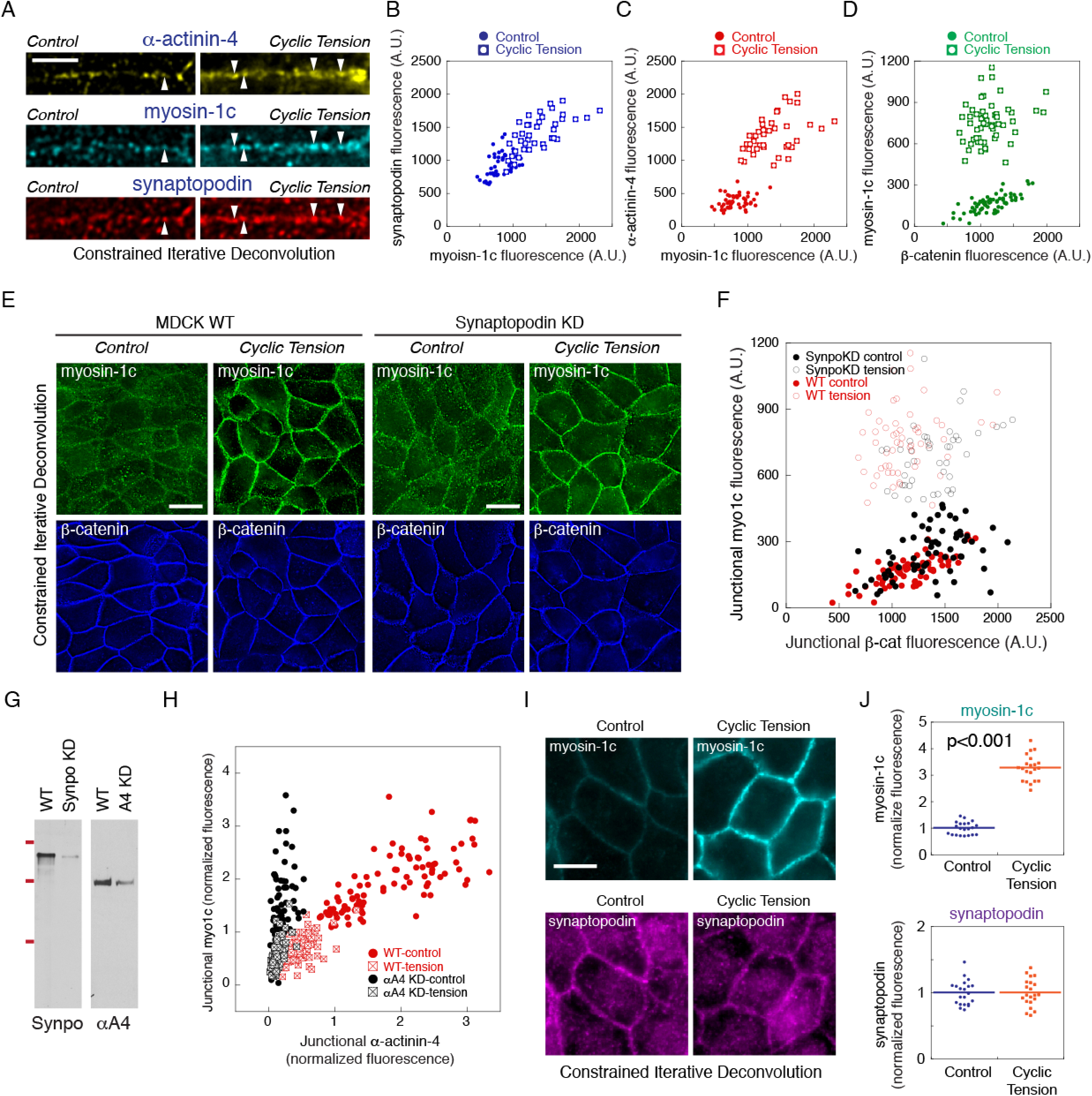
Tension-induced junctional myosin-1c correlates with, but independent of, junctional synaptopodin and *α*-actinin-4. (A) Constrained iterative deconvolution microscopy showing immunofluorescence of myosin-1c, *α*-actinin-4, and synaptopodin in control and cyclic tension treated polarized monolayers. White arrows point to colocalization of myosin-1c, *α*-actinin-4, and synaptopodin. Data is representative of 6 sets of images (N=6) from one experiment out of 4 independent experiments (N=4). (B) Junctional synaptopodin immunofluorescence is plotted against junctional immunofluorescence of myosin-1c. Data is representative of 6 sets of images (N=6) from one experiment out of 4 independent experiments (N=4). (C) Junctional *α*-actinin-4 immunofluorescence is plotted against junctional immunofluorescence of myosin-1c. Data is representative of 6 sets of images (N=6) from one experiment out of 4 independent experiments (N=4). (D) Junctional myosin-1c immunofluorescence is plotted against junctional immunofluorescence of *β*-catenin. Data is representative of 6 sets of images (N=6) from one experiment out of 4 independent experiments (N=4). (E) Constrained iterative deconvolution microscopy showing immunofluorescence of myosin-1c and *β*-catenin in control and cyclic tension treated wild-type (WT) and synaptopodin knockdown (KD) cell monolayers. Data is representative of 12 sets of images (N=12) from one experiment out of 3 independent experiments (N=3). (F) Junctional myosin-1c immunofluorescence is plotted against junctional immunofluorescence of *β*-catenin (*β*-cat) in control (control) and cyclic tension-treated (tension) wild-type (MDCK) and synaptopodin knockdown (synpo KD) cell monolayers. Data is representative of 8 images (N=8) from one experiment out of 3 independent experiments (N=3). (G) Western blots for synaptopodin (Synpo) and *α*-actinin-4 (*α*A4) of total cell lysates from synaptopodin and *α*-actinin-4 knockdown (KD) monolayers. Data is representative of 6 independent experiments (N=6). (H) Junctional myosin-1c (myo1c) immunofluorescence is plotted against junctional immunofluorescence of *α*-actinin-4 (*α*A4) in control (control) and cyclic tension-treated (tension) wild-type (WT) and *α*-actinin-4 knockdown (*α*A4 KD) cell monolayers. Data is representative of 8 images (N=8) from one experiment out of 3 independent experiments (N=3). (I) Constrained iterative deconvolution microscopy showing immunofluorescence of myosin-1c and synaptopodin in control and cyclic tension treated mature cell monolayers. Data is representative of 12 images (N=12) from one experiment out of 4 independent experiments (N=4). (J) Quantification of junctional myosin-1c and synaptopodin immunofluorescence intensities in wild-type (WT) and myosin-1c knockdown (Myo1c KD) monolayers in control and cyclic tension-treated mature cell monolayers. Horizontal lines represent mean fluorescence intensities. Data is representative of 4 independent experiments (N=4).

### Actin-binding function of Myosin-1c is required for α-actinin-4 recruitment

Myosin-1c ATPase defective mutants [42] and a motorless truncation of myosin-1c [43] have previously been used to study its actin-binding and motility function. Here, we generated cell lines expressing EGFP-myosin-1c and several function-defective mutants, including a mutant with defective ATPase activity and no actin binding (EGFP-myosin-1c-R162A) [44-46], a mutant with reduced ATPase activity and weak actin-binding (EGFP-myosin-1c-G389A) [47, 48], a mutant without the motor domain (motorless EGFP-myosin-1c), a mutant with only the motor domain (EGFP-motor only), and a mutant with defective PIP2-binding activity (EGFP-myosin-1c-R903A). We found that EGFP-myosin-1c-R162A and the motorless EGFP-myosin-1c mutant dramatically disrupt the cell-cell interface, causing formation of large blebs on the lateral membrane (Fig. 8A and Supplementary Fig. 12, orange circles). This phenotype is reminiscent of the large blebs seen in myosin-1c knockdown cells (Movies 15-16). Live-cell imaging of the ATPase defective G389A mutant using wide-field microscopy and OS-SIM reveal dynamic and sporadic blebbing at the lateral cell-cell interface (Movies 18-23). These membrane blebbing activities are rarely seen in wild-type cells (Movie 24-25).

**Figure 8.**
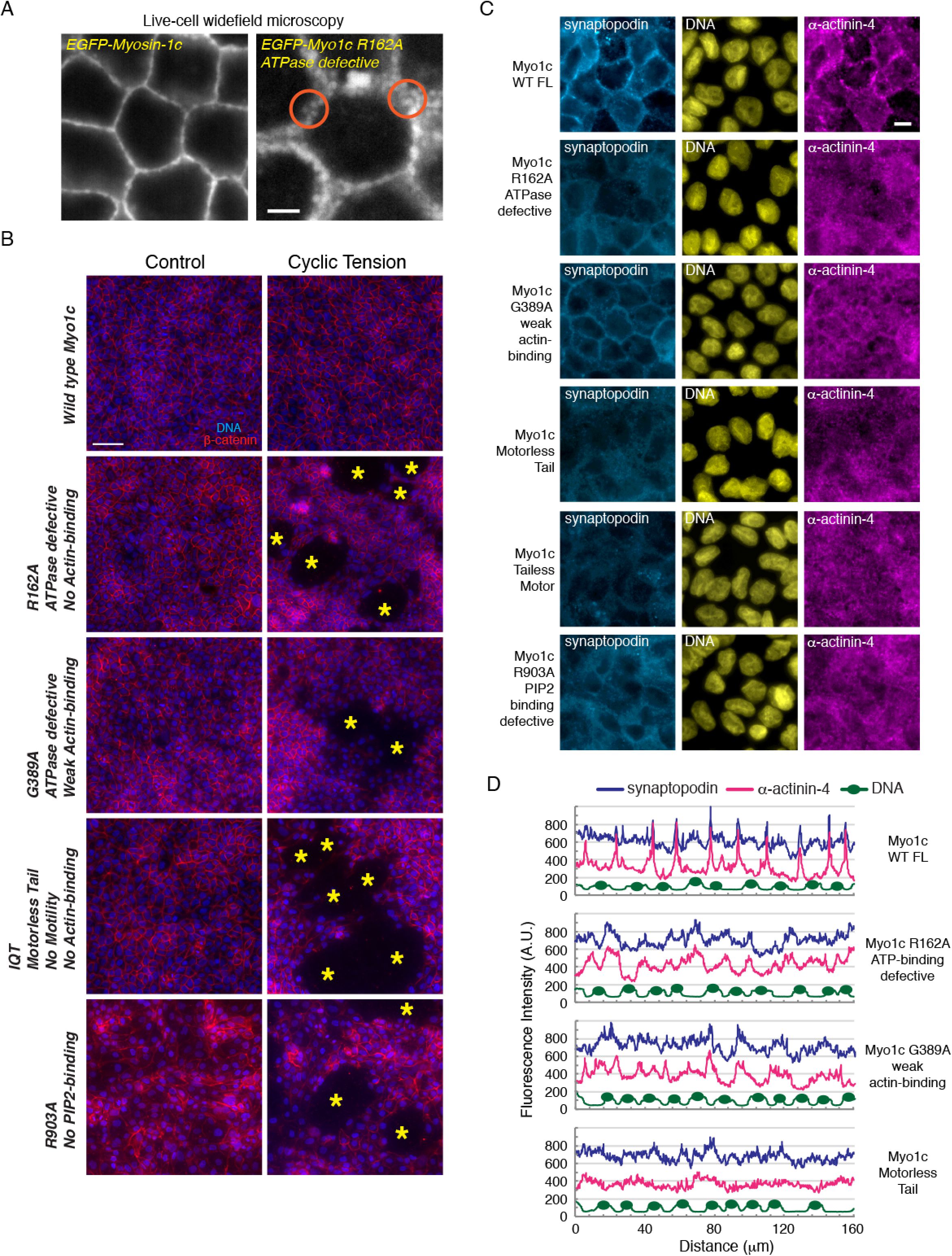
Actin-binding function of Myosin-1c is required for *α*-actinin-4 recruitment. (A) EGFP-myosin-1c-R162A ATPase defective mutant causes bleb formation (orange circles) on the lateral membrane of MDCK cells whereas EGFP-myosin-1c expression does not. Data is representative of 6 sets of live-cell time-lapse movies (N=6). (B) Widefield immunofluorescence images showing *β*-catenin and hoechst stain (DNA) in cell monolayers untreated (control) or treated with 30 min of medium cyclic tension. Cells expressing myosin-1c mutants form intact monolayers under unperturbed condition but became susceptible mechanical stress. Yellow asterisks mark damaged areas that are devoid of cell nuclei. Image set is representative of 12 sets of images (N=12) from one experiment out of 3 independent experiments (N=3). (C) Wide-field immunofluorescence of synaptopodin, *α*-actinin-4, and DNA (hoechst) in monolayers expressing wild-type myosin-1c, myosin-1c-R162A ATPase defective mutant, myosin-1c-G389A weak actin-binding mutant, motorless myosin-1c, myosin-1c motor domain, myosin-1c-R903A PIP2-binding defective mutant. Data is representative of 8 images (N=8) from one experiment out of 3 independent experiments (N=3). Scale bar is 5 μm. (D) Line scan of synaptopodin (blue line), *α*-actinin-4 (pink line, and DNA (Hoechst, green line) levels across several junctions and nuclei (yellow ovals) in monolayers expressing wild-type myosin-1c (Myo1c), myosin-1c-R162A, myosin-1c-G389A, and motorless myosin-1c mutants. Data is representative of 12 line scans (N=12) from one experiment out of 3 independent experiments (N=3).

Expression of myosin-1c mutants in MDCK cells compromises cohesion of the monolayers under mechanical stress (Fig. 8B). When cells expressing myosin-1c mutants were subjected to cyclic intercellular force (∼30 nN per cell at 20 cycles per min for 5 minutes), the monolayers were torn apart, leaving large holes without cells (Fig. 8B, yellow asterisks). These observations are reminiscent of the phenotype of myosin-1c KD, indicating that myosin-1c mutants can act as dominant negatives to weaken intercellular interactions and undermine the integrity of the lateral cell-cell interface.

Expression of EGFP-myosin-1c mutants inhibits the recruitment of synaptopodin and *α*-actinin-4 to the cell junction in mature cell monolayers (Fig. 8C). Line scans across several junctions shows that synaptopodin and *α*-actinin-4 would co-distribute in cells expressing wild-type EGFP-myosin-1c but not in cells expressing EGFP-myosin-1c-R162A, EGFP-myosin-1c-G389A, or the motorless EGFP-myosin-1c mutants (Fig. 8D). These observations indicate that actin binding is necessary for the force transmission function of myosin-1c at the lateral cell-cell interface. Without its actin-binding activity, myosin-1c cannot propagate actomyosin-generated force to cell-cell adhesion and, consequently, is unable to promote tension-dependent recruitment of junctional components. Our current study identifies myosin-1c as the first non-junctional tension-responsive protein on the lateral membrane that is required for the molecular changes associated with adhesion strengthening and junction maturation.

## Discussion

Actomyosin II contractility plays an essential role in adhesion strengthening and the development of cell-cell contacts. Current model describes multiple force-dependent and actin-dependent steps during remodeling of the cell adhesion complex. A repertoire of molecular mechanisms has been discovered including force-induced actin loading onto *α*-catenin, force-induced conformational change in *α*-catenin, force-induced targeting of vinculin, and force-induced targeting of *α*-actinin-4. We had previously shown that synaptopodin and *α*-actinin-4 respond strongly to exogenous applied cyclic force and accumulate at cell-cell interactions to strengthen epithelial cohesion [21]. Here, we discovered a novel tension-sensitive player, myosin-1c, that is upstream and essential for *α*-actinin-4 recruitment during early stages of junction maturation. We advocate that actomyosin II contractility can exert force on cell-cell adhesions through the interaction between cortical actin and the lateral plasma membrane. We demonstrated that myosin-1c is a major determinant of this force transmission pathway and is necessary for tension-dependent junction remodeling. Upon junction maturation, myosin-1c colocalizes with actin-rich junctional complexes on the lateral cell-cell adhesion interface. Our findings offer a complementary mechanism to the established model of mechano-transduction at intercellular interactions and underscore a non-junctional component in the regulation of cell-cell adhesion. We propose that there are at least four phases of junction maturation (Supplementary Fig. 13): 1. increase in cytoplasmic calcium causes global contraction of actomyosin II cytoskeleton, 2. contraction causes inward movement of the plasma membrane that is coupled to cortical actin by myosin-1c, 3. inward movement of the plasma membrane causes cell-cell adhesion to move away from the neighboring cell, 4. movement of adhesions mechanically stimulates tension-dependent junction maturation.

Myosin-1c has been shown to regulate the stability of E-cadherin-based cell-cell contacts by suppressing E-cadherin endocytosis [42]. In a previous study using methanol/acetone fixation, myosin-1c was found to accumulate at cell-cell interface together with E-cadherin. In our current study using paraformaldehyde fixation along with deconvolution microscopy and OS-SIM at high resolution, we found that myosin-1c only overlaps with E-cadherin and *α*-catenin at discreet actin-rich regions of the junction (Fig. 3A). Moreover, myosin-1c strongly overlaps with tension-sensitive synaptopodin and *α*-actinin-4 that are important for adhesion strengthening and junction maturation in epithelial cell sheets [21]. Thus, myosin-1c may not directly inhibit E-cadherin endocytosis but stabilizes the cell junction via a tension-dependent mechanism.

We showed that myosin-1c facilitates the coupling between actin and adhesion complex and plays an essential role in force transmission during actomyosin-II-dependent junction maturation. Knockdown of myosin-1c prevents the accumulation of *α*-actinin-4 at cell-cell contacts without altering its steady-state levels. This effect is similar to our previous study showing that the localization of *α*-actinin-4 can be compromised without changing its cellular level in synaptopodin knockdown cells. Because the recruitment of *α*-actinin-4 is a tension-dependent process, there are at least 2 possibilities that can result in the disruption of its localization to the junction. Possibility one is a disruption of the receptor mechanism at the cell adhesion complex for *α*-actinin-4. Possibility two is a disruption of the force transmission pathway. To distinguish whether the defect is due to *α*-actinin-4 docking or mechano-regulation, we applied exogenous force to cell-cell adhesions in synaptopodin knockdown cell monolayers to test if we can reverse the defect of *α*-actinin-4 localization in synaptopodin knockdown cells. We found that, in the absence of synaptopodin, *α*-actinin-4 no longer responds to intercellular tension. In contrast, we showed that exogenous application of force to myosin-1c-depleted junction fully rescued junctional localization of *α*-actinin-4. Therefore, the receptor mechanism at the junction remains intact in myosin-1c knockdown cells. Because the recruitment of *α*-actinin-4 can be completely reversed by the application of intercellular force, the dependency of *α*-actinin-4 on myosin-1c must be principally biophysical in nature. Once *α*-actinin-4 had been targeted to the junction, its accumulation is maintained independent of myosin-1c. By propagation actomyosin II contractility to cell-cell adhesions, myosin-1c indirectly supports the assembly and organization of junctional components [49, 50], effectively stabilizing the entire intercellular interface.

Myosin-1c can respond to exogenously applied force and accumulate at the cell-cell adhesion interface. However, the mechanism for myosin-1c mechano-regulation is not entirely clear. Knockdown of synaptopodin or *α*-actinin-4 does not affect myosin-1c localization nor its response to intercellular tension, indicating that myosin-1c regulation is independent of synaptopodin and *α*-actinin-4. This uncharacterized upstream mechanism might control myosin-1c targeting through phosphoinositides since myosin-1c localization on the lateral membrane requires its phosphoinositides-binding site [42]. In addition to myosin-1c regulation, PIP2 could directly modulate membrane mechanical properties [51, 52] and indirectly control actomyosin II contractility [53, 54]. Hence, multiple inter-dependent biochemical and biophysical pathways are in place to control the mechanical properties of the lateral membrane and to support tension-driven processes such as cell-cell adhesion and epithelial maturation. We propose that, in epithelial cells, an uncharacterized mechanotransduction mechanism involving lipid metabolism or lipid signalling plays an important role on the lateral membrane to regulate the stiffness of the lateral membrane and the coupling between the plasma membrane and actin cytoskeleton, ultimately the stability and strength of cell-cell adhesion.

After junction maturation, myosin-1c remains sensitive to exogenously applied intercellular force and continues to show strong localization at the lateral membrane. Therefore, myosin-1c is likely to play additional roles in force-dependent processes in epithelial cell sheets. Processes such as collective cell movement and coordinated contraction to shape epithelial cell sheet, could very much depend on proper force transmission between cell-cell adhesions and the membrane-associated actin cortex. By linking the force-generating actomyosin II cytoskeleton to the plasma membrane, myosin-1c positions itself directly in the force-transmission pathways. Thus, myosin-1c is an ideal candidate for force modulation, such as by adjusting its magnitude, the wave-form, or duration. Myosin-1c interacts with actin in a force-sensitive manner and would decrease its detachment rate under load [35]. As a consequence, myosin-1c can hold onto an actin filament longer if the actin filament is pulled. We propose that one possible function of myosin-1c in mature cell monolayers is to provide sustained membrane tension beyond the initial force production step of an actomyosin II contraction. In support of our hypothesis, myosin-1 has been shown to promote the generation of resting membrane tension [55-57], and plays a role in force production [41, 58] and actin compression on the lateral membrane [59, 60]. The interaction of myosin-1c to actin filaments can be fine-tuned by intracellular calcium. Binding of calcium to calmodulin-bound myosin-1c increases the flexibility of the myosin-1c tail and changes its actin gliding properties [38-40, 61, 62]. By regulating myosin-1c through calcium and PIP2, the biophysical characteristics of the force being transmitted, such as the magnitude, duration, or wave-form can be adjusted. In conclusion, myosin-1c is likely to serve multiple functions on the lateral membrane by acting as a linker molecule that possesses the ability to modulate force transmission and adjust tension on the lateral cell-cell interface.

## Materials and Methods

### Antibodies and reagents

Rabbit polyclonal antibodies to synaptopodin were raised against a synthetic peptide corresponding to aa 899-918 of human synaptopodin, SPRAKQAPRPSFSTRNAGIE, that has been coupled to KLH (Pacific immunology). Rabbit polyclonal antibodies to α-actinin-4 were raised in house against a synthetic peptide corresponding to aa 8-24 of human α-actinin-4, NQSYQYGPSSAGNGAGC, that has been coupled to KLH. Rabbit polyclonal antibodies to radixin were raised in house against recombinant full-length human radixin. Antibodies to myosin-1c (sc-136544, mouse), *α*-actinin-4 (sc-49333, goat), *β*-catenin (sc-7963, mouse & sc-7199, rabbit), synaptopodin (sc-50459, rabbit, sc-21536 & sc-21537, goat), *α*-catenin (sc-1495, goat & sc-9988, mouse), p120 (sc-13957, rabbit), vinculin (sc-5573, rabbit), myh9 (sc-98978, rabbit), myh10 (sc-47204, goat) were purchased from Santa Cruz Biotechnology. Rabbit antibodies to myh10 (PRB-445P) were purchased from Biolegend. Antibodies to myosin-1c (ab94539, rabbit) and PIP2 (ab2335, mouse) were purchased from Abcam. Antibodies to E-cadherin (24E1, rabbit) were purchased from Cell Signaling Technology, Inc. Secondary antibodies were purchased from Bio-Rad Laboratories (HRP goat anti–rabbit), Santa Cruz Biotechnology (HRP rabbit anti–goat and goat-anti-mouse), Life Technologies/Invitrogen (FITC and Cy3 goat anti–mouse, FITC and Cy3 goat anti–rabbit, Alexa 488 donkey anti-mouse, Alexa 568 donkey anti-rabbit, Alexa 647 donkey anti-goat). Alexa 647–phalloidin, was purchased from Life Technologies/Invitrogen. Leupeptin, Pefabloc, E-64, antipain, aprotinin, bestatin, and calpain inhibitors I and II were purchased from A.G. Scientific, Inc.

### DNA constructs

ShRNAs for myosin-1c (AACTTCACCAGTGAGGCAGCCTTCATTGA and AATGACAAGAGTGACTGGAAGGTCGTCAG) were synthesized and subcloned into blasticidin-selectable pGFP-B-RS vector (Origene). Knockdowns of *α*-actinin-4 and synaptopodin were performed as published [21]. Briefly, ShRNAs for *α*-actinin-4 (AGGTCCTGTTCCTCTGACTCGGTATCTAT and ACACAGATAGAGAACATCGACGAGGACTT) were synthesized and subcloned into blasticidin-selectable pRS vector and ShRNA for canine synaptopodin (GAGGTGAGATCCAGCACACTTCTGATTGA) was synthesized and subcloned into puromycin-selectable pRS vectors (Origene). Lyn-D3cpV cDNA was kindly provided by Amy Palmer and Roger Tsien [63]. Venus-*α*-actinin-1 cDNA was kindly provided by Fanjie Meng and Frederick Sachs [64]. pcDNA3 EGFP-myosin-1c constructs (EGFP-myosin-1c wild-type full length, EGFP-myosin-1c R162A, EFGP-myosin-1c G389A, EFGP-myosin-1c motorless tail, EFGP-myosin-1c R903A, EGFP-myosin-1c motor domain) were kindly provided by Tokuo H & Lynne Coluccio [42].

### Bait-based crosslinking and mass spectroscopy

Purification of junction-enriched membranes were performed as described [65]. Briefly, frozen rat livers (Pelfrez) were thawed in 2 volumes of 10 mM Hepes, pH8.5/10 mM DTT. Protease inhibitors (see above) were added and the livers were briefly blended in a Waring blender (5 x 15 sec). The liver slush was filtered through 4 layers of cheesecloth to obtain the total liver homogenate. Total liver homogenate was centrifuged at 1000 xg for 30 min. The pellet was homogenized in 10 mM Hepes, pH8.5/10 mM DTT in a dounce homogenizer and centrifuged at 100 xg for 30 min. The supernatant was collected and centrifuged at 1000xg for 30 min. The membrane pellet contains the majority of actin assembly activity and is frozen at -80 degrees before further purification immediately before use. The day of the experiment, membranes were thawed on ice, diluted 1:1 with 10 mM HEPES, pH 8.5 supplemented with 10 mM DTT and homogenized through a 25G needle. The homogenates were spun through a 20% sucrose pad for 10 minutes at 16,000 xg. The supernatant was discarded and the pellet were resuspended with 10 mM HEPES, pH 8.5 supplemented with 10 mM DTT. The homogenate was spun through a 20% sucrose pad for 15 min at 1000 xg. The pellet was discarded and the supernatant was spun through a 20% sucrose pad for 15 min at 16,000 xg. The tiny membrane pellet contains junction-enriched plasma membrane vesicles.

Crosslinking was performed as described [66]. Briefly, a heterobifunctional crosslinker, succinimidyl-diazirine, was used for a 2-step crosslinking reaction via succinimidyl-based chemistry and diazirine-based photochemistry. We covalently attached the crosslinker to recombinant 6-His-tagged *α*-actinin-4 by selectively activating the succinimidyl functional group. The derivatized *α*-actinin-4 was used as “bait” for binding to membranes that had been pre-treated with high salt to remove endogenous *α*-actinin-4. Unbound *α*-actinin-4 was removed by spinning the membranes through a 20% sucrose cushion at 10,000 *g* for 10 min. The membranes were re-suspended in actin assembly buffer and placed under UV to activate the diazirine functional group. The crosslinked *α*-actinin-4 complexes were solubilized in boiling 2% SDS, centrifuged at 16,000 *g* to remove insoluble materials. The supernatant was diluted to a final concentration of 0.2% SDS in 1% TX-100 and incubate overnight with nickel resin(QIAGEN). The nickel resin was washed with 20-bed volumes of 0.5% TX-100 in 20 mM Hepes, pH 7.8 and the crosslinked *α*-actinin-4 complexes were eluted with 5-bed volumes of 0.5% TX-100 in 500 mM imidazole and 20 mM Hepes, pH 7.8. The crosslinker was cleaved by dialysis of the eluate against TCEP for 24 hours at 37^o^C. Cleaved products were separated by SDS-PAGE. Distinct protein bands were excised from Commassie-stained gel and submitted for mass spectroscopy analysis at the University of Illinois Roy J. Carver Biotechnology Center.

### Cell culture & transfection

Madin-Darby canine kidney (MDCK) cells were originally from Kai Simons lab (EMBL, Germany) and a gift from Barry Gumbiner (University of Washington). The cells have been authenticated by staining of E-cadherin and ZO-1 using antibodies that only recognize the canine proteins, RR1 for E-cadherin and R40.76 for ZO-1. The cells are free from mycoplasma contamination as determined by the original source. The cellsMDCK cells were maintained in MEM/Earle’s balanced salt solution supplemented with 25 mM Hepes and 5% FBS. For transfection, cells were incubated in Opti-MEM (Invitrogen) with a 1:1 mixture of DNA/polyethyleimine and selected for 10 days using G418, puromycin, or blasticidin. Clonal cell lines of *α*-actinin-4 and synaptopodin knockdown were obtained as published [21]. Briefly, antibiotic resistant clonal cell lines were expanded and assessed for knockdown efficiency by western blot and immunofluorescence. Clonal cell lines with homogeneous knockdown phenotype were used for a second round of transfection with ShRNA. Secondary clonal cell lines were expanded and assessed for knockdown efficiency by western blot and immunofluorescence. Clonal *α*-actinin-4 knockdown cell lines with knockdown efficiency of >70% were used in this study.

For mixed population of myosin-1c knockdown, antibiotic resistant cells were pooled and used for experiment 14-21 days post-transfection. For homogenously population of myosin-1c knockdown cells, clonal cell lines were expanded and assessed for knockdown efficiency by western blot and immunofluorescence. Antibiotics was used to maintain selection until the cells were ready for experiments.

For live-cell imaging of *α*-actinin-venus (G418 selection), clonal cell lines were expanded and assessed as described [21]. For myosin-1c knockdown in *α*-actinin-venus-expressing cells, stable clonal *α*-actinin-venus cells were transfected with ShRNA for myosin-1c and selected using blasticidin. Clonal myosin-1c knockdown cell lines were expanded and assessed for knockdown efficiency by western blot and immunofluorescence. For live-cell imaging of EGFP-myosin-1c proteins, 24 clonal cell lines expressing EGFP-myosin-1c constructs (EGFP-myosin-1c wild-type, EGFP-myosin-1c R162A, EFGP-myosin-1c G389A, EFGP-myosin-1c motorless tail, EFGP-myosin-1c R903A, EGFP-myosin-1c motor domain) were expanded and assessed for expression efficiency using live-cell microscopy. For live-cell imaging of lynD3cpV, clonal cell lines were expanded and assessed for expression efficiency using live-cell microscopy.

For Live-cell imaging, glass coverslips were soaked in 100% ethanol, and sterilized under UV for 60 min. Sterilized coverslips were coated with 20 ug/ml collagen IV in phosphate buffered saline for 60 min and used immediately for plating of cells.

### Western blot

For comparison of young and mature monolayers, confluent monolayers of wild-type and myosin-1c MDCK cells were trypsinized and replated at high confluent density (10^7^ cells per 10 cm). Cell were allowed to form cell-cell interactions for 2 days (young) or 7 days (mature). Total cell lysates were obtained by solubilizing the cells in SDS-PAGE sample buffer containing 25 mM dithiothreitol, 2% SDS, 50 mM Tris-Cl, 5% glycerol, pH 8.8 and protease inhibitors (10 ug/ml Leupeptin, 1mg/ml Pefabloc, 10 ug/ml E-64, 2 ug/ml antipain, 2 ug/ml aprotinin,50 ug/ml bestatin, 20 ug/ml calpain inhibitors I & 10 ug/ml calpain inhibitor II). Biorad DC detergent compatible protein assay was used to determine total protein concentration in cell lysates. Equal protein amount of cell lysates were used for comparison of junctional protein by western blot.

### Force application in cell monolayer

Cells grown on Transwell-Clear (Corning) were used in tension experiments as described [21]. Briefly, Transwell filter cups were mounted onto a custom-made pressure chamber instrument. Hydraulic pressure was applied to the basal compartment by pumping media into the basal chamber. Pressure was monitor through an outlet from the basal pressure chamber to a pressure gauge. An apical adaptor is mounted onto the apical chamber of the Transwell cup that is filled with cell culture media and sealed with an O-ring. An outlet from the apical adaptor is either exposed to ambient pressure or connected via a bifurcation to a pressure gauge and a syringe hooked up to a syringe pump. Cyclic or pulsatile pressures were applied to cell monolayers by a programmable infuse/withdrawal syringe pump (Lagato SPLG270). All experiments were performed in a 37-degree Celsius room.

### Transepithelial electrical resistance measurement

Cells grown on Transwell-Clear (Corning) were used in transepithelial electrical resistance (TER) measurements. TER were measured using the Epithelial Volt/Ohm meter (EVOM^2^, World Precision Instruments) and chopstick electrodes (STX2, World Precision Instruments). All TER measurements were performed at 37 degrees Celcius.

### Immunofluorescence of cells

Cells grown on Transwell-Clear (Corning) were used in localization studies. For immunofluorescence, cells were rinsed twice in 150 mM NaCl/2 mM CaCl_2_/2 mM MgCl_2_/20 mM Hepes, pH 7.8 and fixed in 1% formaldehyde/150 mM NaCl/2 mM CaCl_2_/2 mM MgCl_2_/20 mM Hepes, pH 7.8 at 4°C for 2 hours. The reaction was quenched with Tris in quenching buffer (0.05% Triton X-100/50 mM Tris/100 mM NaCl/20 mM Hepes, pH 8.0) for 3 hours. After rinsing in staining buffer (0.01% Triton X-100/100 mM NaCl/20 mM Hepes, pH 7.5), the cells were incubated with primary antibodies in staining buffer overnight. After rinsing in staining buffer three times, the cells were incubated in secondary antibodies in staining buffer for 90 min. Then, the cells were rinsed three times in staining buffer and incubated with fluorescently labeled phalloidin or Hoechst for 60 min. Finally, the cells were rinsed three times in staining buffer and post-stain fixed with 1% formaldehyde in staining buffer for 60 min. Transwell filters were excised using a razor blade and mounted on glass slides with ProLong Diamond antifade (Invitrogen).

### Image acquisition

For figures 1-6, 9, S1, S3-4, S6-7, S9-14, S15A, S16-17, images were collected in 200-nm steps using Axio Imager.Z2m microscope equipped with Apotome.2 (Zeiss) and X-cite 120 LED (Lumen Dynamics). For Optical Sectioning Structured Illumination Microscopy (OS-SIM), 7 phases/images were collected per each constructed image using an alpha Plan-Apochromat 100x/1.46 oil DIC M27 objective (Zeiss) and Apotome.2. For OS-SIM, images were collected using a 4K ORCA-Flash4.0 V2 digital cmos camera (ORCA-ER; Hamamatsu Photonics). For figures 9, S14-14, 17-19, low magnification wide field images were collected using a Plan Apochromat 20X/0.8 objective (Zeiss) or a FLUAR 40X/1.3 oil objective (Zeiss) with a 2K Optimos digital cmos camera (Qimaging).

For figures 3 and S9-10, wide field images were collected in 200-nm steps using Axio Imager.Z2m microscope, the ORCA-Flash4.0 V2 camera, and the alpha Plan-Apochromat 100x/1.46 oil DIC M27 objective. Wide field optical z images were deconvolved using the Zen2 pro deconvolution module (nearest neighbor, fast iterative, or constrained iterative algorithms as stated in figure legends).

For figures 6-7 and S15C, images were collected in 200-nm steps using an inverted microscope (IX-71; Olympus), a 1K charge-coupled device camera (Cool SNAp HQ, Applied Precision), a 60X/1.42 oil objective with a 1.6X auxiliary magnification, and SoftWorx DMS software (Applied Precision). Wide field optical z images were deconvolved using the enhanced ratio constrained iterative deconvolution algorithm with 10 iteration cycles (Applied Precision).

### Live-cell imaging

Cells grown on collagen-coated glass coverslips were mounted up-side-down onto an in-house fabricated glass slide chamber. The chamber was assembled by attaching a 1.5x1.5 cm medical-grade silicone gel (PediFix) onto a sterilized glass slide. The center of the silicone gel was excised and used as a sink for media during imaging. Live-cell imaging was performed in FluoroBrite/DMEM (Gibco) media containing 1% fetal bovine serum and 10 mM HEPES, pH 7.5. Sample temperature was maintained at 35 degrees Celsius on a heated stage and an objective heater (PeCon) mounted onto the Axio Imager.Z2m microscope.

For live-cell imaging of *α*-actinin-venus and EGFP-myosin-1c, stable clonal cell lines in wild-type and myosin-1c knockdown background were used. Stable clonal cell lines expressing EGFP-myosin-1c wild-type and mutant proteins (EGFP-myosin-1c, EGFP-myosin-1c R162A, EFGP-myosin-1c G389A, EFGP-myosin-1c motorless tail, EFGP-myosin-1c R903A, EGFP-myosin-1c motor domain) were used in live-cell imaging. Live-cell wide field images were collected using a FLUAR 40X/1.3 oil objective (Zeiss) and the ORCA-Flash4.0 V2 camera. Live-cell OS-SIM, 5 phases/images were collected per each constructed image using an alpha Plan-Apochromat 100x/1.46 oil DIC M27 objective (Zeiss), Apotome.2 (Zeiss), and a 2K Optimos camera (Qimaging) mounted onto an Axio Imager.Z2m microscope (Zeiss).

### Calcium Imaging

Intracellular calcium was monitored using 2 calcium sensors: a lipid-modifiable calcium sensor protein, lynD3cpV (CFP-cpVenus FRET pair) and a calcium-binding small molecule, Rhod-4. Changes in calcium is monitored by excited emission of lynD3cpV (excitation of CFP and emission of cpVenus with YFP filter) and cpVenus (excitation of cpVenus and emission of cpVenus with YFP filter). For figure 2C & D, images were collected with a FLUAR 40X/1.3 oil objective (Zeiss) using a Axio Imager.Z2m microscope and X-cite 120 LED (Lumen Dynamics). For OS-SIM in figure 2E, 5 phases/image were collected per each constructed image using Axio Imager.Z2m microscope equipped with Apotome.2 (Zeiss) and X-cite 120 LED (Lumen Dynamics). For calcium sensitive Rhod-4, cells were loaded according to the manufacturer’s instruction (ab112157, ABCAM). Briefly, cells were incubated in Rhod-4 dye-loading solution for 1 hour and image immediately with a FLUAR 40X/1.3 oil objective (Zeiss) using a Axio Imager.Z2m microscope and X-cite 120 LED (Lumen Dynamics).

### Image Analysis

All images were corrected for chromatic shift (using fluorescent beads as fiducial marks) on the X, Y, Z-axes for each fluorescence channel before were used for analysis. Quantitation of immunofluorescence intensity was performed in ImageJ. All measured intensities were subtracted from background signal (an area with no cells) before being used for statistical analyses and calculation of intensity ratios. Each junctional region is outlined manually with a freehand drawing tool. The mean pixel intensity of each defined junctional region is used for comparison of junction localization of individual junctional protein. For calculation of Pearson’s correlation coefficient R, intensities of individual pixel within the defined junctional region were used and each pixel corresponds to 45 x 45 nm of the imaged sample. Line intensity graphs were generated in Excel (Microsoft) using pixel intensities from original images. Quantitation of Rhod-4 fluorescence intensity and measurements of cell diameter, cell area, and perimeter were performed using Zen2 pro analysis tools (Zeiss).

For measurement of cell width, a line tool was use to draw lines from one side of the cell to the other. The position of the line was determined empirically by moving and rotating the line 360 degrees so that the line is as perpendicular to the junctions as possible.

### Image Processing

For figure generation, images were cropped, contrasted, and scaled using Photoshop software (Adobe) before importing into Illustrator (Adobe). For movie generation, individual images of cropped cells were imported into QuickTime (Apple) to generate movie files. Composite images were generated using ImageJ (NIH) or Photoshop (Adobe).

### Data Analysis and Statistics

All experiments had been repeated at least three times. At least 6 data set from each experiment were collected. All p-values were calculated using non-paired student t-tests. Linear regression was fitted using original data points. All analyses were performed using KaleidaGraph software (Synergy).

### Data Availability

All data are available from the corresponding author.

## Acknowledgements

I thank Nivetha Kannan for the generation of cell lines, immunofluorescence staining, data organization and analysis. I thank Lydia Lee for propagating cell lines. I thank Kevin Huang for western blotting. I thank Nilmani Singh for DNA work.

## Supplementary Figure Legends

**Supplementary Figure 1. Myosin-1c is chemically crosslinked in a mature junctional complex with *α*-actinin-4.** Myosin-1c peptides identified by mass spectroscopy (red letters) from a proximity bait-based crosslinking experiment using *α*-actinin-4 as bait (see Methods).

**Supplementary Figure 2. Myosin-1c co-distributes with synaptopodin and *α*-actinin-4 at linear junctions and junctional vertices along the lateral membrane in polarized epithelial monolayer.** Linear regression analysis was performed for each protein pair. Pearson’s correlation coefficient (R) between each protein-pair was calculated from 500-2000 pixels per protein per z-image per one set of data. Data set is representative of 6 sets of data (N=6) from one experiment out of 3 independent experiments (N=3).

**Supplementary Figure 3. Myosin-1c overlaps with *α*-catenin and E-cadherin at discreet regions along the lateral junctions in polarized epithelial monolayer.** (A) Optical sectioning structured illumination microscopy showing staining of myosin-1c (M1c), *α*-catenin (Acat), and E-cadherin (Ecad) at the apical junction (Z=0 μm), sub-apical junction (Z=1.6 μm), lateral junctions (Z=3.2 and 4.8 μm). DNA is stained with Hoechst. For each Z-image, the top panel shows the X-Z, Y-Z, and X-Y images and the bottom panel shows a representative linear junction from the X-Y-image. Orange scale bar is 10 μm and white scale bar is 2 μm. Data is representative of 10 sets of images (N=10) from one experiment out of 3 independent experiments (N=3). (B) Merge images of the representative linear junction shown in A. White arrows point to colocalization of myosin-1c, *α*-catenin, and E-cadherin. White arrows point to colocalization of myosin-1c, *α*-catenin, and E-cadherin. Orange arrows point to colocalization of myosin-1c and *α*-catenin. Red arrows point to colocalization of *α*-catenin and E-cadherin. Pearson’s correlation coefficient (R) between 2 proteins at the linear junction. Correlations were calculated from 1000-1500 pixels per protein per z-image per one set of data (see Figure S8). Data set is representative of 6 sets of data (N=6) from one experiment out of 3 independent experiments (N=3). Scale bar is 2 μm.

**Supplementary Figure 4. Myosin-1c weakly correlates with *α*-catenin and E-cadherin along the lateral junctions in polarized epithelial monolayer.** Linear regression analysis was performed for each protein pair. Pearson’s correlation coefficient (R) between each protein-pair was calculated from 500-1500 pixels per protein per z-image per one set of data. Data set is representative of 6 sets of data (N=6) from one experiment out of 3 independent experiments (N=3).

**Supplementary Figure 5. Synaptopodin colocalizes with myosin IIB and actin at the apical junction in polarized epithelial monolayer.** (A) Pixel intensity of synaptopodin, myosin IIB, and actin (phalloidin) staining showing overlapping peaks at junctions (i-v). (B) Optical sectioning structured illumination microscopy showing staining of myosin-1c, actin (phalloidin), and myosin IIB at the apical junction. For each Z-image, the top panel shows the X-Z, Y-Z, and X-Y images and the bottom panel shows a representative junction area from the X-Y-image. Yellow arrows point to puncta with synaptopodin, myosin IIB, and actin (phalloidin). Image is representative of 10 images (N=10) from one experiment out of 3 independent experiments (N=3). Yellow scale bar is 5 μm and white scale bar is 1 μm.

**Supplementary Figure 6. Myosin-1c strongly co-distributes with actin on the lateral junctions in polarized epithelial monolayer.** (A) Optical sectioning structured illumination microscopy showing staining of myosin-1c (M1c), actin (phalloidin, Phal), and myosin IIB (MIIB) at the apical junction (Z=0 μm), sub-apical junction (Z=1 μm), lateral junctions (Z=2.6 and 4.2 μm), and basal junction (Z=6 μm). DNA is stained with Hoechst. For each Z-image, the top panel shows the X-Z, Y-Z, and X-Y images and the bottom panel shows a representative linear junction from the X-Y-image, except for the basal Z=6 μm which shows the area with stress fibers. Data is representative of 10 sets of images (N=10) from one experiment out of 3 independent experiments (N=3). Orange scale bar is 10 μm and white scale bar is 2 μm. (B) Merge images of the representative linear junction shown in A. Pearson’s correlation coefficient (R) between 2 proteins at the linear junction. Correlations were calculated from 1000-2000 pixels per protein per z-image per one set of data (see Figure S5). Data set is representative of 6 sets of data (N=6) from one experiment out of 3 independent experiments (N=3). Scale bar is 2 μm.

**Supplementary Figure 7. Myosin-1c correlates with actin on the lateral junctions.** Linear regression analysis was performed for each protein pair. Pearson’s correlation coefficient (R) between each protein-pair was calculated from 500-1500 pixels per protein per z-image per one set of data. Data set is representative of 6 sets of data (N=6) from one experiment out of 3 independent experiments (N=3).

**Supplementary Figure 8. Deconvolution and wide field images of myosin-1c, *β*-catenin, and actin staining in heterogeneous myosin-1c knockdown cell monolayer, corresponding to image in figure 2.** A stack of z-images collected at 0.2 μm intervals were deconvolved using nearest neighbor, fast iterative, and constrained iterative algorithms. Data is representative of 16 sets of images (N=16) from one experiment out of 3 independent experiments (N=3). Scale bar is 10 μm.

**Supplementary Figure 9. Whole cell contraction is preceded by calcium spike and characterized by inward movement of the lateral plasma membrane.** (A) Measurements of cell width and changes in cytoplasmic calcium using a lipid-modified intramolecular FRET sensor, lynD3cpV from Movie 3. Orange dotted lines represent beginning of a calcium spike. Orange solid line represents the beginning of a contraction. (B) Time-lapse frames from Movie 5 showing transient increase in calcium followed by contraction. LynD3cpV FRET (excitation of donor and emission of acceptor), cpVenus (excitation of acceptor and emission of acceptor contraction) and merge images of FRET and cpVenus are shown. Pink lines mark the original cell width of the cell being stretched at a corner by its neighboring cell. Arrowheads mark the junctional region of the cell being stretched. Calcium spike is indicated by a change in FRET signal in the cell being stretched. Data is representative of 6 sets of live-cell time-lapse movies (N=6).

**Supplementary Figure 10. Epithelial contractions associate with junction accumulation of *α*-actinin.** (A) Time-lapse frames from Movie 6 showing transient increase in the accumulation of venus-*α*-actinin at the cell junction during a junctional contraction (blue asterisk) in a mature epithelial monolayer. Blue (J1) and orange (J2) circles mark the cell junctional areas associated with the contracting (D1) and non-contracting (D2) junction, respectively. Blue and orange lines delineate the cell diameters positioned at a contracting (D1) and a non-contracting (D2) junction in the same cell. Data is representative of 6 sets of live-cell time-lapse movies (N=6). Scale bar is 10 μm. (B) Measurement of cell width at a contracting (Distance 1) and a non-contracting (Distance 2) junction in A and Movie 6. Pink solid line marks the frames shown in A. Pink dotted line marks the drop and recovery of the cell diameter. (C) Measurement of junctional intensity of venus-*α*-actinin at a contracting (Junction 1) and a non-contracting (Junction 2) area corresponding to the junctions in A, B and movie 8. Pink solid line marks the frames shown in A. Pink dotted line shows the corresponding rise and recovery of junctional venus-*α*-actinin intensity. (D) Time-lapse frames from Movie 7 showing transient decrease in cell width during a whole cell contraction (red and yellow asterisks) in a maturing monolayer. Yellow line marks the cell width. Data is representative of 6 sets of live-cell time-lapse movies (N=6). Scale bar is 10 μm. (E) Measurements of cell width of the contracting cell shown in D and Movie 7. Left and right insets show junctional venus-*α*-actinin at t=0 min and t=120 min, respectively. Scale bar is 2 μm. (F) Quantitation of junctional venus-*α*-actinin of the contracting cell shown in D-E at t=0 and t=4.5 hr. Bar graph shows the mean and standard errors of 6 separate junctional regions of the contraction cell (N=6). Data is representative of 6 sets of live-cell time-lapse movies (N=6). (G) Time-lapse frames from movie 11 showing whole cell contraction (green and red asterisks) along the lateral membrane (apical Z=5 μm to basal Z=0 μm) in a young monolayer with no junction accumulation of *α*-actinin (green and yellow arrows). Pink arrows point to venus-*α*-actinin on the nuclear envelope. Data is representative of 12 sets of live-cell time-lapse movies (N=12). Scale bar is 10 μm. (H) Measurements of the cell shown in G and movie 11. Apical junctional level of venus-*α*-actinin (Z=4 μm) gradually increases after contraction (purple asterisk). (I) Measurements of the cell shown in G and movie 11. Lateral junctional level of venus-*α*-actinin (Z=1 μm) transiently decreases during contraction (green asterisk). (J) Measurements of the cell shown in G and movie 11. Nuclear level of venus-*α*-actinin (Z=1 μm) transiently increases during contraction (red asterisk).

**Supplementary Figure 11. Time-lapse frames from Movie 12 showing transient increase in the accumulation of venus-*α*-actinin on the nuclear envelope during whole cell contraction (from apical Z= 6 μm to basal Z=0 μm).** Yellow, pink, green, orange, and blue arrows mark transient accumulation of venus-*α*-actinin on the nuclear envelopes in cells A, B1, B2, C1, and C2, respectively. Data is representative of 12 sets of live-cell time-lapse movies (N=12). Scale bar is 10 μm.

**Supplementary Figure 12. Myosin-1c mutations defective in actin-binding or motor function cause bleb formation (orange circles) at the lateral membrane in polarized cell monolayers.** Widefield images showing expression of EGFP-myosin-1c, EGFP-myosin-1c-R162A ATPase defective mutant, EGFP-myosin-1c-G389A weak actin-binding mutant, EGFP-myosin-1c motorless mutant, EGFP-motor domain, and EGFP-myosin-1c-R903A PIP2-binding defective mutant in polarized cell monolayers. Image set is representative of 8 sets of images (N=8) from one experiment out of 3 independent experiments (N=3).

**Supplementary Figure 13. Interactive mechano-regulation of junction maturation - a working model.** We propose that there are at least four phases of junction maturation: 1. increase in cytoplasmic calcium causes global contraction of actomyosin II cytoskeleton, 2. contraction causes inward movement of the plasma membrane that is coupled to cortical actin by myosin-1c, 3. inward movement of the plasma membrane causes cell-cell adhesion to move away from the neighboring cell, 4. movement of adhesions mechanically stimulates tension-dependent junction maturation.

## Movie Legends

**Movie 1.** MDCK monolayer expressing EGFP-myosin-1c at 2-day post-confluency. The movie shows 2 contractions as indicated by contraction along the lateral cell-cell interface (apical Z= 6 μm to basal Z= 0 μm).

**Movie 2.** MDCK monolayer expressing EGFP-myosin-1c at 2-day post-confluency showing multiple short contractions.

**Movie 3.** MDCK monolayer expressing lynD3cpV at 2-day post-confluency showing calcium spikes preceding cellular contractions. Increase in FRET is not associated with increase in YFP.

**Movie 4.** Live-OS-SIM of MDCK monolayer expressing lynD3cpV at 2-day post-confluency showing inward movement of plasma membrane during cellular contraction.

**Movie 5.** MDCK monolayer expressing lynD3cpV at 4-day post-confluency showing calcium spikes preceding cellular contractions. Increase in FRET is not associated with increase in YFP.

**Movie 6.** MDCK monolayer expressing venus-*α*-actinin at 5-day post-confluency showing a regional contraction associated with an increase in venus-*α*-actinin fluorescence at the junction.

**Movie 7.** MDCK monolayer expressing venus-*α*-actinin at 5-day post-confluency showing repeated contractions and increase in junctional accumulation of venus-*α*-actinin.

**Movie 8.** MDCK monolayer expressing venus-*α*-actinin at 4-day post-confluency showing many whole cell contractions. The contractions are associated with constriction of the cell along the entire lateral Z-axis (apical Z= 8 μm to basal X= 0 μm). Contractions correlate with transient accumulation of venus-*α*-actinin on the nuclear envelope.

**Movie 9.** MDCK monolayer expressing venus-*α*-actinin at 1-day post-confluency showing one single whole cell contraction. The contraction is associated with constriction of the cell along the entire lateral Z-axis (apical Z= 6 μm to basal X= 0 μm). Contraction correlates with a transient accumulation of venus-*α*-actinin on the nuclear envelope.

**Movie 10.** MDCK monolayer expressing venus-*α*-actinin at 2-day post-confluency showing several whole cell contractions. The contractions are associated with constriction of the cell along the entire lateral Z-axis (apical Z= 9 μm to basal X= 0 μm). Contractions correlate with transient accumulation of venus-*α*-actinin on the nuclear envelope.

**Movie 11.** MDCK monolayer expressing venus-*α*-actinin at 1-day post-confluency showing one single whole cell contraction. The contraction is associated with constriction of the cell along the entire lateral Z-axis (apical Z= 5 μm to basal X= 0 μm). Contraction correlates with a transient accumulation of venus-*α*-actinin on the nuclear envelope.

**Movie 12.** MDCK monolayer expressing venus-*α*-actinin at 1-day post-confluency showing asynchronous cell contractions in a group of cells. The contractions are associated with constriction of the cell along the entire lateral Z-axis (apical Z= 6 μm to basal X= 0 μm). Contractions correlate with transient accumulation of venus-*α*-actinin on the nuclear envelope.

**Movie 13.** Myosin-1c knockdown cells expressing venus-*α*-actinin at 7-day post-confluency showing one single cell contraction. The contraction is associated with transient accumulation of venus-*α*-actinin on the nuclear envelope.

**Movie 14.** Myosin-1c knockdown cells expressing venus-*α*-actinin at 7-day post-confluency showing one single cell contraction. The contraction is associated with transient cell expansion and accumulation of venus-*α*-actinin on the nuclear envelope.

**Movie 15.** Myosin-1c knockdown cells expressing venus-*α*-actinin at 7-day post-confluency showing lack of venus-*α*-actinin accumulation at cell junction and constant formation of large blebs at cell-cell interface.

**Movie 16.** Myosin-1c knockdown cells expressing venus-*α*-actinin at 7-day post-confluency showing lack of venus-*α*-actinin accumulation at cell junction and constant formation of large blebs at cell-cell interface.

**Movie 17.** MDCK monolayer expressing venus-*α*-actinin at 7-day post-confluency showing accumulation of venus-*α*-actinin at cell junction. Venus-*α*-actinin is dynamic at the junction but no large blebs are formed.

**Movie 18.** EGFP-Myosin-1c-G389A expressing cells at 7-day post-confluency showing dynamic blebbing at cell-cell interface.

**Movie 19.** EGFP-Myosin-1c-G389A expressing cells at 7-day post-confluency showing dynamic blebbing at cell-cell interface.

**Movie 20.** Live-OS-SIM showing EGFP-Myosin-1c-G389A expressing cells at 7-day post-confluency with dynamic blebbing at cell-cell interface.

**Movie 21.** Live-OS-SIM showing EGFP-Myosin-1c-G389A expressing cells at 7-day post-confluency with dynamic blebbing at cell-cell interface.

**Movie 22.** Live-OS-SIM showing EGFP-Myosin-1c-G389A expressing cells at 7-day post-confluency with dynamic blebbing at cell-cell interface.

**Movie 23.** Live-OS-SIM showing EGFP-Myosin-1c-G389A expressing cells at 7-day post-confluency with dynamic blebbing at cell-cell interface.

**Movie 24.** EGFP-Myosin-1c expressing cells at 7-day post-confluency showing a lack of constant blebbing at cell-cell interface.

**Movie 25.** Live-OS-SIM showing EGFP-Myosin-1c expressing cells at 7-day post-confluency showing a lack of constant blebbing at cell-cell interface.

